# Transmembrane helices are an overlooked and evolutionarily conserved source of major histocompatibility complex class I and II epitopes

**DOI:** 10.1101/2021.05.02.441235

**Authors:** Richèl J.C. Bilderbeek, Maxim Baranov, Geert van den Bogaart, Frans Bianchi

## Abstract

Cytolytic T cell responses are predicted to be biased towards membrane proteins. The peptide-binding grooves of most haplotypes of histocompatibility complex class I (MHC-I) are relatively hydrophobic, therefore peptide fragments derived from human transmembrane helices (TMHs) are predicted to be presented more often as would be expected based on their abundance in the proteome. However, the physiological reason of why membrane proteins might be over-presented is unclear. In this study, we show that the over-presentation of TMH-derived peptides is general, as it is predicted for bacteria and viruses and for both MHCI and MHC-II. Moreover, we show that TMHs are evolutionarily more conserved, because single nucleotide polymorphisms (SNPs) are present relatively less frequently in TMH-coding chromosomal regions compared to regions coding for extracellular and cytoplasmic protein regions. Thus, our findings suggest that both cytolytic and helper T cells respond more to membrane proteins, because these are evolutionary more conserved. We speculate that TMHs therefore are less prone to escape mutations that enable pathogens to evade T cell responses.

## 1 Introduction

Our immune system fights diseases and infections from pathogens, such as fungi, bacteria or viruses. An important part of the acquired immune response, that develops specialized and more specific recognition of pathogens than the innate immune response, are T cells which recognize peptides, called epitopes, derived from antigenic proteins presented on Major Histocompatibility Complexes (MHC) class I and II on the cell surface.

The MHC proteins are heterodimeric complexes encoded by the HLA (Human Leukocyte Antigens) genes. In humans, the peptide binding groove of MHC-I is made by only the alpha subunit. There are three classical forms of MHC-I, hallmarked by a highly polymorphic alpha chain called HLA-A, HLA-B and HLA-C, that all present epitopes to cytolytic T cells. For MHC-II, both the alpha and the beta chains contribute to the peptide binding groove. There are three classical forms of MHC-II as well, called HLA-DR, HLA-DQ and HLA-DP, that all present epitopes to helper T cells. Each MHC complex can present a subset of all possible peptides. For example, HLA-A and HLA-B have no overlap in which epitopes they bind [1]. Moreover, the HLA genes of humans are highly polymorphic, with hundreds to thousands of different alleles, and each different HLA allele is called an MHC haplotype and presents a different subset of peptides [2].

Humans mostly express two haplotypes per MHC form, one from the parental and one from the maternal chromosome, and therefore an individual’s immune system detects only a fraction of all possible peptide fragments. However, at the population level, the coverage of pathogenic peptides that are detected is very high, because of the highly polymorphic MHC genes. It is therefore believed that MHC polymorphism improves immunity at the population level, as mutations in a protein that disrupt a particular MHC presentation at the individual level, so-called escape mutations, will not affect MHC presentation for all haplotypes present in the population [3].

Many studies are aimed at identifying the repertoire of epitopes that are presented in any MHC haplotype and determining which epitopes will result in an immune response, as this will for instance aid the design of vaccines. These studies have led to the development of prediction algorithms that allow for very reliable *in silico* predictions of the binding affinities of peptides [4, 5, 6]. For example, [6] found that, of the 432 peptides that were predicted to bind to MHC, 86% were experimentally confirmed to do so.

Using these prediction algorithms, we recently predicted that peptides derived from transmembrane helices (TMHs) will be more frequently presented by MHC-I than expected based on their abundance [7]. Moreover, we showed that some well-known immunodominant peptides stem from TMHs. This overpresentation is attributed to the fact that the peptide-binding groove of most MHC-I haplotypes is relatively hydrophobic, and therefore hydrophobic TMH-derived peptides have a higher affinity to bind than their soluble hydrophilic counterparts.

TMHs are hydrophobic as they need to span the hydrophobic lipid bilayer of cellular membranes. They consist of an alpha helix of, on average, 23 amino acids in length. TMHs can also be predicted with high accuracy from a protein sequence by bioinformatics approaches [8, 9, 10, 11, 12, 13], for example, [11] found that, from 184 transmembrane proteins (TMPs) with known topology, 80% of the TMH predictions replicated this finding.

TMHs are common structures in the proteins of humans and microbes. Different TMH prediction tools estimate that 15-39% of all proteins in the human proteome contain at least one TMH [14]. However, the physiological reason why peptides derived from TMHs would be presented more often than peptides stemming from soluble (i.e., extracellular or cytoplasmic) protein regions is unknown. We hypothesized that the presentation of TMH residues is evolutionary selected for, because TMHs are less prone to undergo escape mutations. One reason to expect such a reduced variability (and hence evolutionary conservation) in TMHs, is that these are restricted in their evolution by the functional requirement to span a lipid bilayer. Due to this requirement, many of the amino acids genuinely present in TMHs are limited to the ones with hydrophobic side chains [15, 16]. Therefore, we speculated that the TMHs of pathogens might have a lower chance to develop escape mutations, as many mutations will result in a dysfunctional TMH and render the protein inactive.

This study had two objectives. First, we aimed to generalize our findings by predicting the presentation of peptides from different kingdoms of life and for both MHC-I and -II. From these *in silico* predictions, we conclude that TMH-derived epitopes are presented more often than expected by chance, in a human, viral and bacterial proteome, and for most haplotypes of both MHC-I and II. We confirmed the presentation of TMH-derived peptides by re-analysis of peptide elution studies. Second, we tested our hypothesis that TMHs are more evolutionary conserved than soluble protein regions. Our analysis of human single nucleotide polymorphisms (SNPs) showed that random point mutations are indeed less likely to occur within TMHs. These findings strengthen the emerging notion that TMHs are important for the T cell branch of the adaptive immune system, and hence are of overlooked importance in vaccine development.

## 2 Methods

### 2.1 Predicting TMH epitopes

To predict how frequently epitopes overlapping with TMHs are presented, a similar analysis strategy was applied as described in [7] for several haplotypes of both MHC-I and MHC-II, and for a human, viral and bacterial proteome. To summarize, for each proteome, all possible 9-mers (for MHC-I) or 14-mers (MHC-II) were derived. For each of these peptides, we determined if it overlapped with a predicted TMH and if it was predicted to bind to each haplotype.

For MHC-I, 9-mers were used, as this is the length most frequently presented in MHC-I and was used in our earlier study [7]. For MHC-II, 14-mers were used, as these are the most frequently occurring epitope length [17]. A human (UniProt ID UP000005640 9606), viral (SARS-CoV-2, UniProt ID UP000464024) and bacterial (*Mycobacterium tuberculosis*, UniProt ID UP000001584) reference proteome was used. TMHMM [8] was used to predict the topology of the proteins within these proteomes. To predict the affinity of an epitope to a certain MHC haplotype, EpitopePrediction [7] for MHC-I and MHCnuggets [18] for MHC-II was used. The 13 MHC-I haplotypes used in this study are the same as used in the previous study [7]. For MHC-II, haplotypes were selected with a phenotypic frequency of at least 14% in the human population [19], resulting in 21 MHC-II haplotypes.

In previous work, it was found that the over-presentation of TMH-derived peptides can be explained from the hydrophobicity of the MHC-I binding cleft [7]. Here, a similar analysis was applied, by correlating the percentage of predicted TMH-derived epitopes versus the mean hydrophobicity of all peptides.

This study differs in one important aspect from our previous work [7]. The definition of a binder differs from [7]: in the current study, a peptide is called a binder if, for a certain haplotype, any of its 9-mer or 14-mer peptides have an IC50 value in the lowest 2% of all peptides within a *proteome* (see supplementary Tables 4 and 5 for values), whereas the previous study defined a binder as having an IC50 in the lowest 2% of the peptides within a *protein*. This revised definition precludes bias of proteins that give rise to no or only very few MHC epitopes. To verify that the results are similar, a side by side comparison was performed shown in the supplementary materials.

### 2.2 Peptide elution studies

To obtain experimental evidence that epitopes derived from TMHs are presented in MHC, peptide elution studies for MHC-I [20] and MHC-II [17] were reanalyzed. For each of the detected epitopes, its possible location(s) in a human reference proteome, with UniProt ID UP000005640 9606, was mapped. For the epitopes that were present in the proteome exactly once, the topology of the proteins in which these epitopes were located was predicted using both TMHMM [8] and PureseqTM [13]. From this topology, we determined if the epitope overlapped with a TMH.

The full analysis can be found at https://github.com/richelbilderbeek/bbbq_article_issue_157.

#### 2.2.1 Evolutionary conservation of TMHs

To determine the evolutionary conservation of TMHs, human single nucleotide polymorphisms (SNPs) were first collected that resulted in a single amino acid substitution, and we then determined if this substitution occurred within a predicted TMH or not.

As a data source, multiple NCBI (https://www.ncbi.nlm.nih.gov/) databases were used: the *dbSNP* [21] database, which contains 650 million catalogued non-redundant humane variations (called RefSNPs, https://www.ncbi.nlm.nih.gov/snp/docs/RefSNP_about/), and the databases *gene* (for gene names, [22]) and *protein* (for proteins sequences, [23]).

The first query was a call to the *gene* database for the term ’membrane protein’ (in all fields) for the organism *Homo sapiens*. This resulted in 1,077 gene IDs (on December 2020). The next query was a call to the *gene* database to obtain the gene names from the gene IDs. Per gene name, the *dbSNP* NCBI database was queried for variations associated with the gene name. As the NCBI API constrains its users to three calls per second (to assure fair use), we had to limit the extent of our analysis.

The number of SNPs was limited to the first 250 variations per gene, resulting in *≈*61k variations. Only variations that result in a SNP for a single amino acid substitution were analyzed, resulting in *≈*38k SNPs. The exact amounts can be found in the supplementary materials, Tables 9 and 10.

SNPs were picked based on ID number, which is linked to their discovery date. To verify that these ID numbers are unrelated to SNP positions, the relative positions of all analyzed SNPs in a protein were determined. This analysis showed no positional bias of the SNPs, as shown in supplementary figure 15.

Per SNP, the *protein* NCBI database was queried for the protein sequence. For each protein sequence, the protein topology was determined using PureseqTM. Using these predicted protein topologies, the SNPs were scored to be located within or outside TMHs.

## 3 Results

### 3.1 TMH-derived peptides are predicted to be over-presented in MHC-I

Figure 1A shows the predicted presentation of TMH-derived peptides in MHCI, for a human, viral and bacterial proteome. Per MHC-I haplotype, it shows the percentage of binders that overlap with a TMH with at least one residue. The horizontal line shows the expected percentage of TMH-derived epitopes that would be presented, if TMH-derived epitopes would be presented just as likely as epitopes derived from soluble regions. For 11 out of 13 MHC-I haplotypes, TMH-derived epitopes are predicted to be presented more often than the null expectation, for a human and bacterial proteome. For the viral proteome, 12 out of 13 haplotypes present TMH-derived epitopes more often than expected by chance. The extent of the over-presentation between the different haplotypes is similar for the probed proteomes, which strengthens our previous conclusion [7] that the hydrophobicity of the MHC-binding groove is the main factor responsible for the predicted over-presentation of TMH-derived peptides.

**Figure 1:**
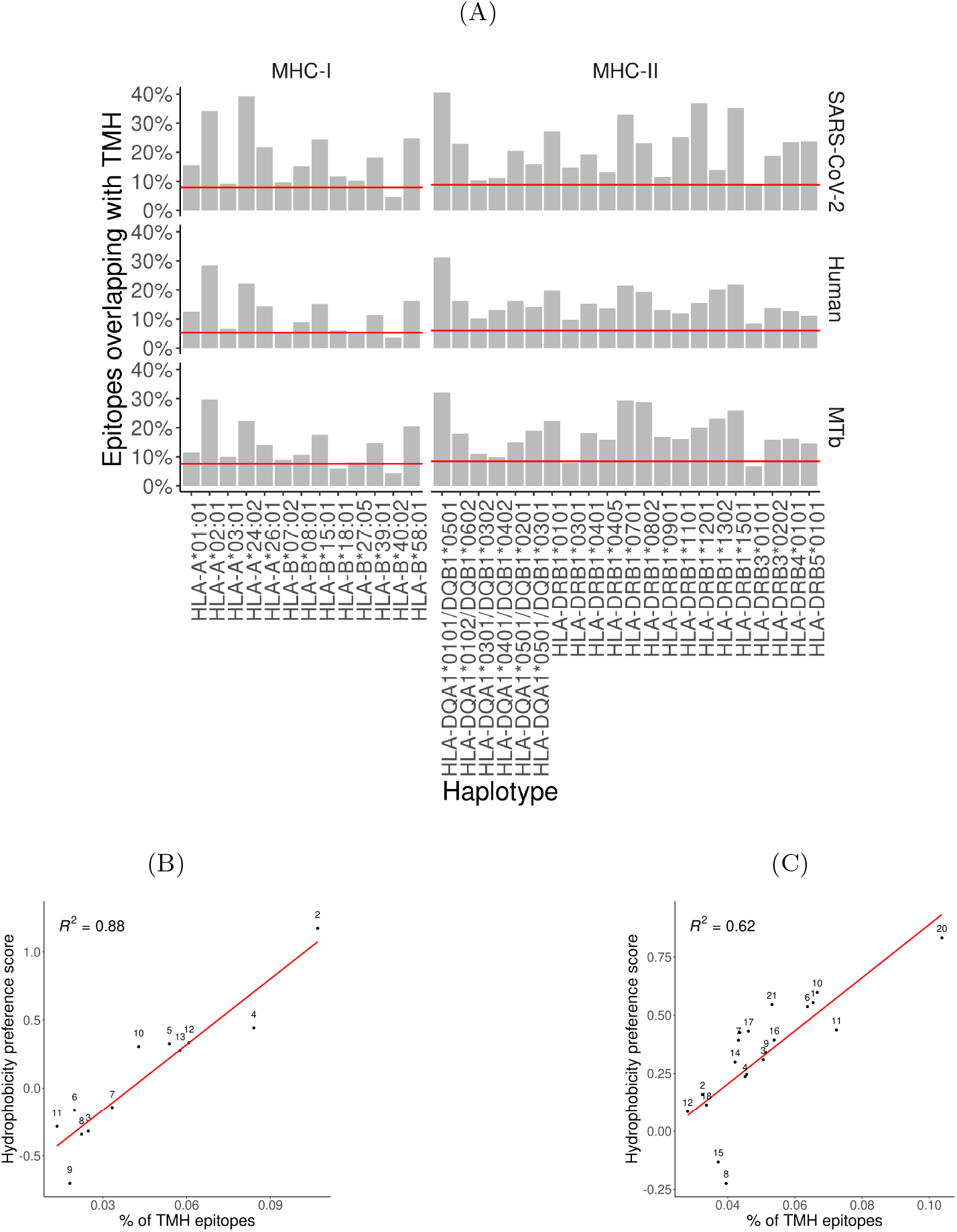
Over-presentation of TMH-derived epitopes on most MHC-I and -II haplotypes. **(A)** The percentage of epitopes for MHC-I and -II haplotypes that are predicted to overlap with TMHs for the proteomes of SARS-CoV-2 (top row), human (middle row) and *M. tuberculosis* (bottom row). The red lines indicate the percentages as expected by chance. See supplementary Tables 7 and 8 for the exact TMH and epitope counts. **(B-C)** Correlation between the percentages of predicted TMH-derived epitopes and the hydrophobicity score of all predicted epitopes for MHC-I **(B)** and MHC-II haplotypes **(C)**. Red curve: linear regression analysis. Labels are shorthand for the HLA haplotypes, see the supplementary Table 6 for the names.

### 3.2 TMH-derived peptides are predicted to be over-presented in MHC-II

We next wondered if the over-representation of TMH-derived peptides would also be present for MHC-II. Figure 1A shows the percentages of MHC-II epitopes predicted to be overlapping with TMHs for our human, viral and bacterial proteomes. We found that TMH-derived peptides are over-presented in all of the 21 MHC-II haplotypes, for a human, bacterial and viral proteome, except for HLA-DRB3*0101 in *M. tuberculosis*. See supplementary Table 8 for the exact TMH and epitope counts.

### 3.3 The over-presentation of TMH-derived peptides is caused by the hydrophobicity of the MHC peptide binding groove

For MHC-I, we previously showed that the over-presentation of TMH-derived peptides is caused by the hydrophobicity of the peptide binding grooves [7]. Figures 1B and 1C show the extent of over-presentation of TMH-derived epitopes as a function of the hydrophobicity preference score for the different haplotypes. An assumed linear correlation explains 88% of the variability in MHC-I. For MHC-II, 62% of the variability is explained by hydrophobicity. This strengthens our previous finding [7] and indicates that TMH-derived peptides are over-presented because the peptide binding grooves of most MHC-I and -II haplotypes are relatively hydrophobic.

### 3.4 Experimental validation of presentation of TMH-derived peptides

To obtain experimental confirmation that peptides stemming from TMHs are presented in MHC-I and MHC-II, two peptide elution studies were reanalyzed: For MHC-I, peptides presented *in vivo* by the (humane) haplotypes HLA-A and B were sequenced [20], for MHC-II these were haplotypes DQ2.5, DQ2.2, and DQ7.5 [17]. Figure 2 shows the percentages of epitopes derived from TMHs found in the MHC-I and MHC-II elution studies, for the two topology prediction tools TMHMM [8] and PureseqTM [13]. Regardless of the prediction tool, at least 100 epitopes were predicted to be derived from a TMH for each condition. From these findings, it is robustly predicted that epitopes derived from TMHs are presented in both MHC-I and MHC-II. See the supplementary Table 3 for the exact values.

**Figure 2:**
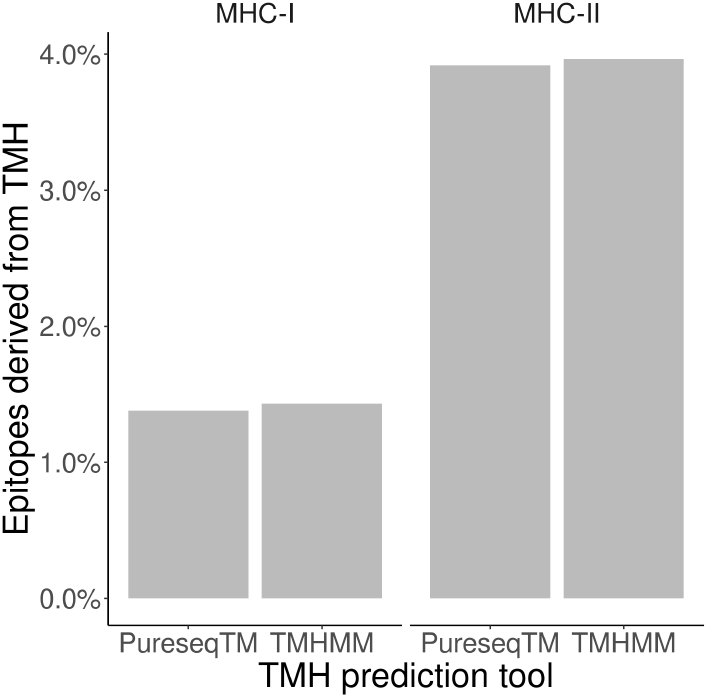
Robust prediction that TMH epitopes are presented *in vivo*. Bars show the percentage of peptides obtained from elution studies that is derived from a TMH, for MHC-I and -II, for two TMH prediction tools.

### 3.5 Human TMHs are evolutionarily conserved

We addressed the question whether there is an evolutionary advantage in presenting TMHs. We determined the conservation of TMHs by comparing the occurrences of SNPs located in TMHs or soluble protein regions for the genes coding for membrane proteins. We obtained 911 unique gene names associated with the phrase ’membrane protein’, which are genes coding for both membrane-associated proteins (MAPs, which have no TMH) and transmembrane proteins (TMPs, which have at least one TMH). These genes are linked to 4,780 protein isoforms, of which 2,553 are predicted to be TMPs and 2,237 proteins are predicted to be MAPs. We obtained 37,630 unique variations, of which 9,621 are SNPs that resulted in a straightforward amino acids substitution, of which 6,062 were located in predicted TMPs. See supplementary Tables 9 and 10 for the detailed numbers and distributions of SNPs.

Per protein, we calculated two percentages: (1) the percentage of the total protein predicted to be TMHs, and (2) the percentage of SNPs located within these predicted TMHs. Each percentage pair was plotted in figure 3B. The proportion of SNPs found in TMHs varied from none (i.e. all SNPs were in soluble regions) to all. To determine if SNPs were randomly distributed over the protein, we performed a linear regression analysis, and added a 95% confidence interval on this regression. This linear fit nearly goes through the origin and has a slope below the line of equality, which shows that less SNPs are found in TMHs than expected by chance.

**Figure 3:**
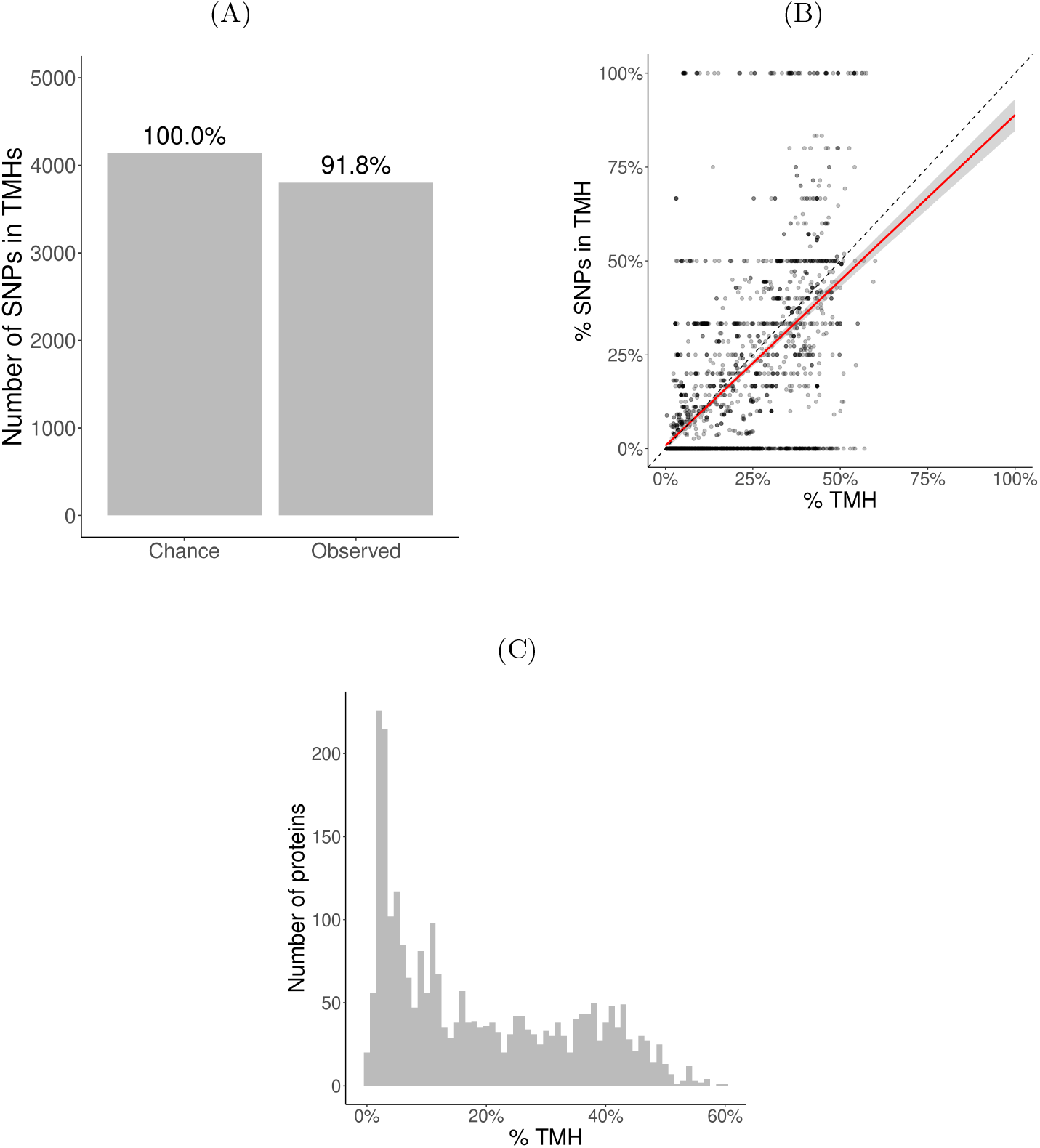
Evolutionary conservation of human TMHs. **(A)** The number of SNPs in TMHs as expected by chance (left bar) and found in the dbSNP database (right bar). Percentages show the relative conservation of SNPs in TMHs found. **(B)** Percentage of SNPs found in TMHs. Each point shows for one protein the predicted percentage of TMH (*x*-axis) and the observed occurrence of SNPs being located within a TMH (*y*-axis). The dashed diagonal line shows the line of equality (i.e., equal conservation of TMHs and soluble protein regions). The red line indicates a linear fit, the gray area its 95% confidence interval. **(C)** Distribution of the percentages of TMH in the TMPs used in this study.

We determined the probability to find the observed amount of SNPs in TMHs by chance, i.e., when assuming SNPs occur just as likely in soluble domains as in TMHs. We used a binomial Poisson distribution, where the number of trails (*n*) equals the number of SNPs, which is 21,208. The probability of success for the *i*th TMP (*p_i*), is the percentage of residues within a TMH per TMP. These percentages are shown as a histogram in figure 3C. The expected number of SNPs expected to be found in TMHs by chance equals Σ *p ≈* 4, 141. As we observed 3,803 SNPs in TMHs, we calculated the probability of having that amount or less successes. We used the type I error cut-off value of *α* = 2.5%. The chance to find, within TMHs, this amount or less SNPs equals 6.8208*·*10*^−^*^11^. We determined the relevance of this finding, by calculating how much less SNPs are found in TMHs, when compared to soluble regions, which is the ratio between the number of SNPs found in TMHs versus the number of SNPs as expected by chance. In effect, per 1000 SNPs found in soluble protein domains, one finds 918 SNPs in TMHs, as depicted in figure 3A.

We split this analysis for TMPs containing only a single TMH (so-called single-membrane spanners) and TMPs containing multiple TMHs (multi-membrane spanners). We hypothesized that single-membrane spanners are less conserved than multi-membrane spanners, because multi-membrane spanners might have protein-protein interactions between their TMHs, for example to accommodate active sites, and thus might have additional structural constraints. From the split data, we did the same analysis as for the total TMPs. Figure 4C shows the percentages of TMHs for individual proteins as a function of the percentage of SNPs located in TMHs. For both single- and multi-spanners, a linear regression shows that less SNPs are found in TMHs, than expected by chance.

**Figure 4:**
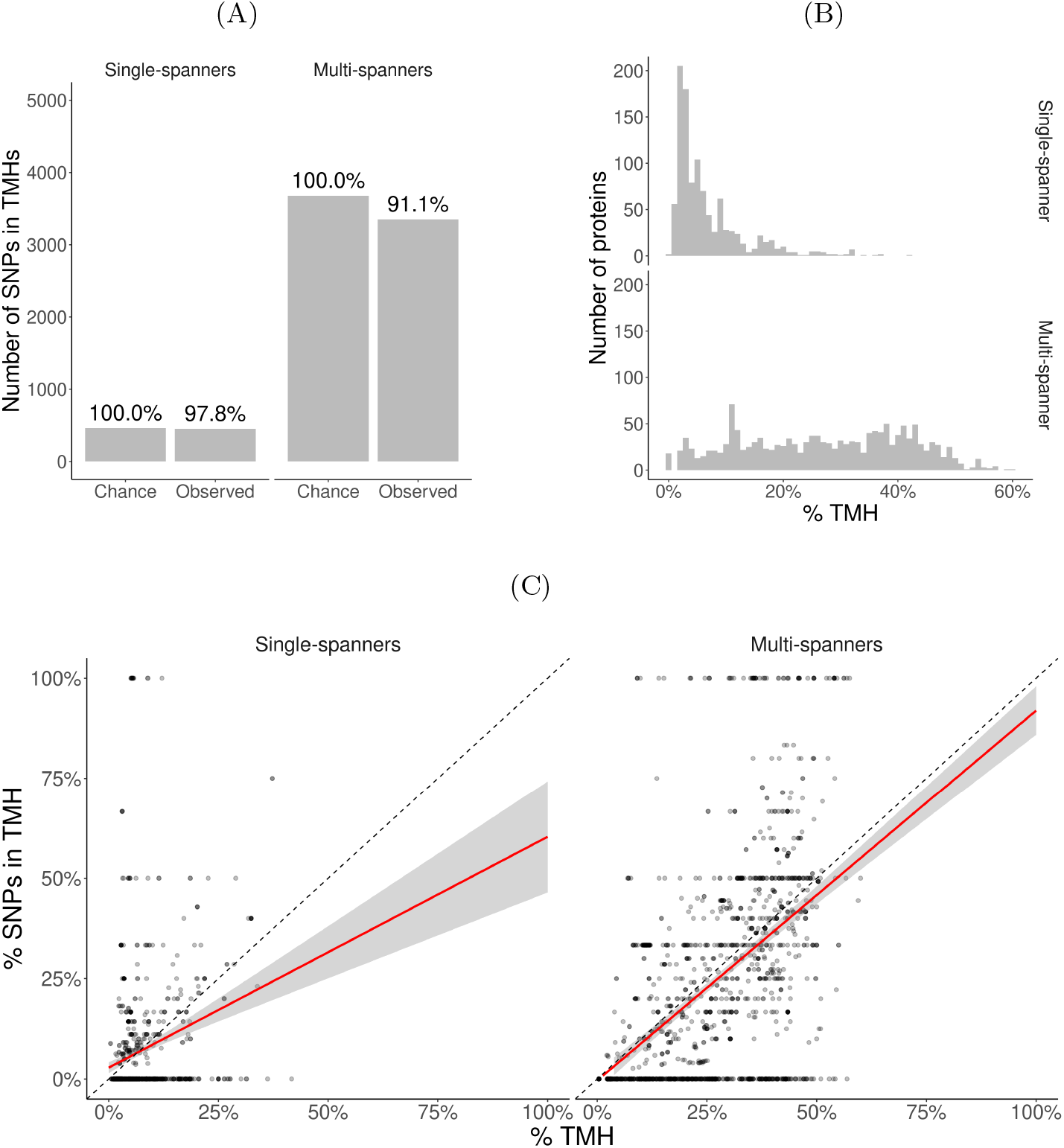
Membrane proteins with multiple TMHs are evolutionary more conserved than proteins with only a single TMH. **(A)** The number of SNPs in TMHs as expected by chance and observed in the dbSNP database, for TMPs with one TMH (single-spanners) and multiple TMHs (multi-spanners). Percentages show the relative conservation of SNPs in TMHs found. **(B)** Distribution of the proportion of amino acids residing in the plasma membrane. **(C)** Percentage of SNPs found in TMPs predicted to have only a single (left) or multiple (right) TMHs. Each point shows for one protein the predicted percentage of TMH (*x*-axis) and the observed occurrence of SNPs being located within a TMH (*y*-axis). The dashed diagonal lines show the line of equality (i.e., equal conservation of TMHs and soluble protein regions). The red line indicates a linear fit, the gray area its 95% confidence interval.

We also determined the probability to find the observed amount of SNPs by chance in single- and multi-spanners. For single-spanners, we found 452 SNPs in TMH, where *≈* 462 were expected by chance. The chance to observe this or a lower number by chance is 0.319. As this chance was higher than our *α* = 0.025, we consider this no significant effect. For the multi-spanners, we found 3,351 SNPs in TMH, where *≈* 3, 678 were expected by chance. The chance to observe this or a lower number by chance is 8.315841 *·* 10*^−^*^12^, which means this number is significantly less as explained by variation.

Also, for single- and multi-spanners, we determined the relevance of this finding by calculating how much less SNPs are found in TMHs when compared to soluble regions, as depicted in figure 4A. In effect, per 1,000 SNPs found in soluble protein domains, one finds 978 SNPs in TMHs of single-spanners and 911 SNPs in TMHs of multi-spanners.

## 4 Discussion

Epitope prediction is important to understand the immune system and for the design of vaccines. In this study, we provide evidence that epitopes derived from TMHs are a major but overlooked source of MHC epitopes. Our bioinformatics predictions indicate that TMH-derived epitopes are presented to both cytolytic and helper T cells more often than expected by chance, regardless of the organism. Moreover, reanalysis of peptide elution studies confirmed the presentation of TMH-derived epitopes. Finally, our SNP analysis shows that TMHs are evolutionary more conserved than solvent-exposed protein regions.

### 4.1 Mechanism of MHC presentation of TMH-derived epitopes

Although our data show that TMH-derived epitopes are presented in MHC-I and MHC-II, the molecular mechanisms of how integral membrane proteins are processed for MHC presentation are largely unknown [7]. Most prominently, the fundamental principles of how TMHs are extracted from their hydrophobic lipid environments into the aqueous vacuolar lumen, and their prior or subsequent proteolytic processing are unresolved.

A first possibility is that the extraction of TMPs from the membrane is mediated by the ER-associated degradation (ERAD) machinery. For MHC class I (MHC-I) antigen presentation of soluble proteins, the loading of the epitope primarily occurs at the endoplasmatic reticulum (ER). The chaperones tapasin (TAPBP), ERp57 (PDIA3), and calreticulin (CALR) [24] first assemble and stabilize the heavy and light chains of MHC-I. Later, this complex binds to the transporter associated with antigen processing (TAP) leading to the formation of the so-called peptide-loading complex (PLC). The PLC drives import of peptides into the ER and mediates their subsequent loading into the peptide-binding groove of MHC-I [25]. Membrane proteins first will have to be extracted from the membrane before they become amenable to this MHC-I loading by the PLC. In the ER, this process can be orchestrated by the ERAD machinery, consisting of several chaperones that recognize TMPs, ubiquitinate them, and extract them from the ER membrane into the cytosol (retrotranslocation) for proteasomal degradation [26, 27]. Similar to the peptides generated from soluble proteins, the TMP-derived peptides might then be re-imported by TAP into the ER for MHC-I loading. This ERAD-driven antigen retrotranslocation might be facilitated by lipid bodies (LBs) [28], since LBs can serve as cytosolic sites for ubiquitination of ER-derived cargo [29].

A second possibility is that TMPs are proteolytically processed by intramembrane proteases that cleave TMHs while they are still membrane embedded. Supporting this hypothesis is the well established notion that peptides generated by signal peptide peptidases (SPPs), an important class of intramembrane proteases that cleave TMH-like signal sequences, are presented on a specialized class of MHC-I called HLA-E [30]. The loading of peptides generated by SPP onto MHC-I does not depend on the proteosome and TAP, possibly because the peptides are directly released into the lumen of the ER [30]. However, this mechanism would not explain how multispanner polytopic membrane proteins can be processed for antigen presentation, because SPPs only cleave TMH-like signal sequences at the N-terminus of a protein. Nevertheless, the presentation of peptides with a high hydrophobicity index was shown to be independent of TAP as well [31], suggesting the TMH peptides might perhaps be released directly in the ER lumen by other intramembrane proteases.

A third possibility is that peptide processing and MHC-loading occur in multivesicular bodies (MVBs) [30]. TMPs can be routed from the plasma membrane and other organelles by vesicular trafficking to endosomes. Eventually, these TMPs can be sorted by the endosomal sorting complexes required for transport (ESCRT) pathway into luminal invaginations that pinch off from the limiting membrane and form intraluminal vesicles. This thus results in MVBs where the membrane proteins destined for degradation are located in intraluminal vesicles. Upon the fusion of MVBs with lysosomes, the entire intraluminal vesicles including the TMPs are degraded [32]. Via this mechanism, TMPs might well be processed for antigen presentation, particularly since the loading of MHC class II molecules is well understood to occur in MVBs [33, 34, 35]. However, such processing of membrane proteins in MVBs for antigen presentation poses a problem, because complexes of HLA-DR with its antigen-loading chaperon HLA-DM were only observed on intraluminal vesicles, but not on the limiting membranes of MVBs [35], indicating that epitope loading of MHC-II also occurs at intraluminal vesicles. This observation hence raises the question how the intraluminal vesicles carrying the TMPs destined for antigen presentation can be selectively degraded, while the intraluminal vesicles carrying the MHC-II remain intact. A second problem is that phagosomes carrying internalized microbes lack intraluminal vesicles, and it is hence unclear how TMPs from these microbes would be routed to MVBs for MHC-II loading [35].

Alternatively to the enzymatic degradation of lipids in MVBs by lipases [36, 37], they might be oxidatively degraded by reactions with radical oxygen species (ROS) produced by the NADPH oxidase NOX2 [38]. This oxidation can result in a destabilization and disruption of membranes [38] and might thereby lead to the extraction of TMPs. Due to the hydrophobic nature of TMHs, however, the extracted proteins will likely aggregate and it is unclear how these aggregates would be processed further for MHC loading.

### 4.2 T cells recognize different protein regions than B cells

An important implication from the over-presentation of TMH-derived epitopes is that T cells will largely recognize different protein regions than B cells. Presentation of antigens by MHC-II is important for the activation of naive B cells by helper T cells. For this activation, B cells first ingest antigen that is bound to their B cell receptor, and subsequently present peptides derived from this antigen in MHC-II to helper T cells. Following their activation by the T cells, B cells mature into plasma cells and release antibodies which recognize the same part of the antigen as the original B cell receptor. B cell receptors and antibodies will thus recognize solvent-exposed regions of antigens that are accessible for binding to the B cell receptor. However, the results from our study predict that most MHC-II haplotypes present relatively hydrophobic peptides, which are less likely to be solvent-exposed. It is unknown why B and T cells seem to predominantly recognize different protein regions, but one possibility might be that this lowers the chance of B cell mediated autoimmune diseases, because auto-reactive B and T cells recognizing different parts of the same antigen would need to be present for breakage of B cell tolerance.

### 4.3 Evolutionary conservation of TMHs

In general, one might expect that evolutionary selection results in an immune system that as most attentive for protein regions that are essential for the survival, proliferation and/or virulence or pathogenic microbes, as these will be most conserved. In SARS-CoV-2, for example, there is preliminary evidence that the strongest selection pressure is upon residues that change its virulence [39]. These regions, however, may only account for a small part of a pathogen’s proteome. Additionally, the structure and function of these essential regions might differ widely between different pathogenic proteins. Because of this scarcity and variance in targets, one can imagine that it will be mostly unfeasible to provide innate immune responses against such rare essential protein regions, as suggested in a study on influenza [40], where it was found that the selection pressure exerted by the immune system was either weak or absent.

Evolutionary selection of pathogens by a host’s immune system, however, is likelier to occur for proteomic patterns that are general, over patterns that are rare. While essential catalytic sites in a pathogenic proteome might be relatively rare, TMHs are common and thus might be a more feasible target for evolution to respond to. Indeed, we have found the signature of evolution when both factors, that is, TMHs and catalytic sites are likely to co-occur, which is in TMPs that span the membrane at least twice. In contrast to single-spanners, where we found no significant evolutionary conservation, the TMHs of multi-spanners are more evolutionary conserved than soluble protein regions. Likely, the TMHs in many multi-spanners need to interact which each other for correct protein structure and function and they might hence be more structurally constrained compared to the TMHs of single-spanners. Thus, we speculate that the human immune system is more attentive towards TMHs in multi-spanners, as these are evolutionarily more conserved.

There have been more efforts to assess the conservation of TMHs, using different methodologies. One such example is [41], in which aligned protein sequence data was used. Also this study found that TMHs are evolutionarily more conserved, as the mean amino acid substitution rate in TMHs is about ten percent lower, which is a similar value as we found. Another example is a study that estimated the conservation scores for TMHs and soluble regions based on alignments of evolutionary related proteins, and also found that TMHs are more conserved, with a conservation score that was 17% higher in TMHs [42]. Note that the last study also found that mutations in human TMHs are likelier to cause a disease, in line with our conclusion that TMHs are more conserved.

Together, from this study, two important conclusions can be drawn. First, the MHC over-presentation of TMHs is likely a general feature and predicted to occur for most haplotypes of both MHC-I and -II and for humans as well as bacterial and viral pathogens. Second, TMHs are genuinely more evolutionary conserved than soluble protein motifs, at least in the human proteome.

## Abbreviations

MAP: Membrane-associated protein
TMH: Transmembrane helix
TMP: Transmembrane protein

## 5 Acknowledgments

We thank the Center for Information Technology of the University of Groningen for its support and for providing access to the Peregrine high performance computing cluster. FB is funded by a Veni grant from the Netherlands Organization for Scientific Research (016.Veni.192.026) and an Off-Road Grant from the Dutch Medical Science Foundation (ZonMW 04510011910005). GvdB is funded by a Young Investigator Grant from the Human Frontier Science Program (HFSP; RGY0080/2018), and a Vidi grant from the Netherlands Organization for Scientific Research (NWO-ALW VIDI 864.14.001). GvdB has received funding from the European Research Council (ERC) under the European Union’s Horizon 2020 research and innovation programme (grant agreement No. 862137.

## 6 Data Accessibility

All code is archived at http://github.com/richelbilderbeek/someplace, with DOI https://doi.org/12.3456/zenodo.1234567.

## 7 Authors’ contributions

RJCB and FB conceived the idea for this research. MVB helped with the proteome analysis of *M. tuberculosis*. RJCB wrote the code. RJCB, MB, GvdB and FB wrote the article.

## A Supplementary materials

### A.1 Differences with Bianchi et al., 2017

A part of this study does the same analysis as Bianchi et al., 2017. mainly concern the use of different software and a different definition of what an MHC binder is.

The earlier study defined a peptide an MHC binder if *within the protein* in which it was found, is was among the peptides with the 2% lowest IC50 values. This can be seen at https://github.com/richelbilderbeek/bianchi_et_al_2017/blob/master/predict-binders.R, where the binders are written to file.

However, in this study, an MHC binder is defined as a peptide within a *proteome* in which it is found, that is among the peptides with the 2% lowest IC50 values. Subsection A.8 shows the IC50 values for a binder per haplotype. We believe that our revised definition is more correct, as it overcomes bias from proteins with very low numbers of peptides and/or MHC-predicted binders.

Our previous study used the TMHMM web server to predict TMHs. The desktop version of TMHMM, however, gives an error message on the 25 seleno-proteins found in the human reference proteome. For the sake of reproducible research, we used the desktop version (as we can call it from scripts) and, due to this, we removed the selenoproteins from this analysis.

To verify if the previous and the current method give rise to notable difference, we show a side-by-side comparison in figures 5A and 5B. The figures that haplotypes that over-present or under-present TMH-derived epitopes, do so in both studies. The extent to which TMH-derived epitopes are presented, however, is more extreme in our current setup.

**Figure 5:**
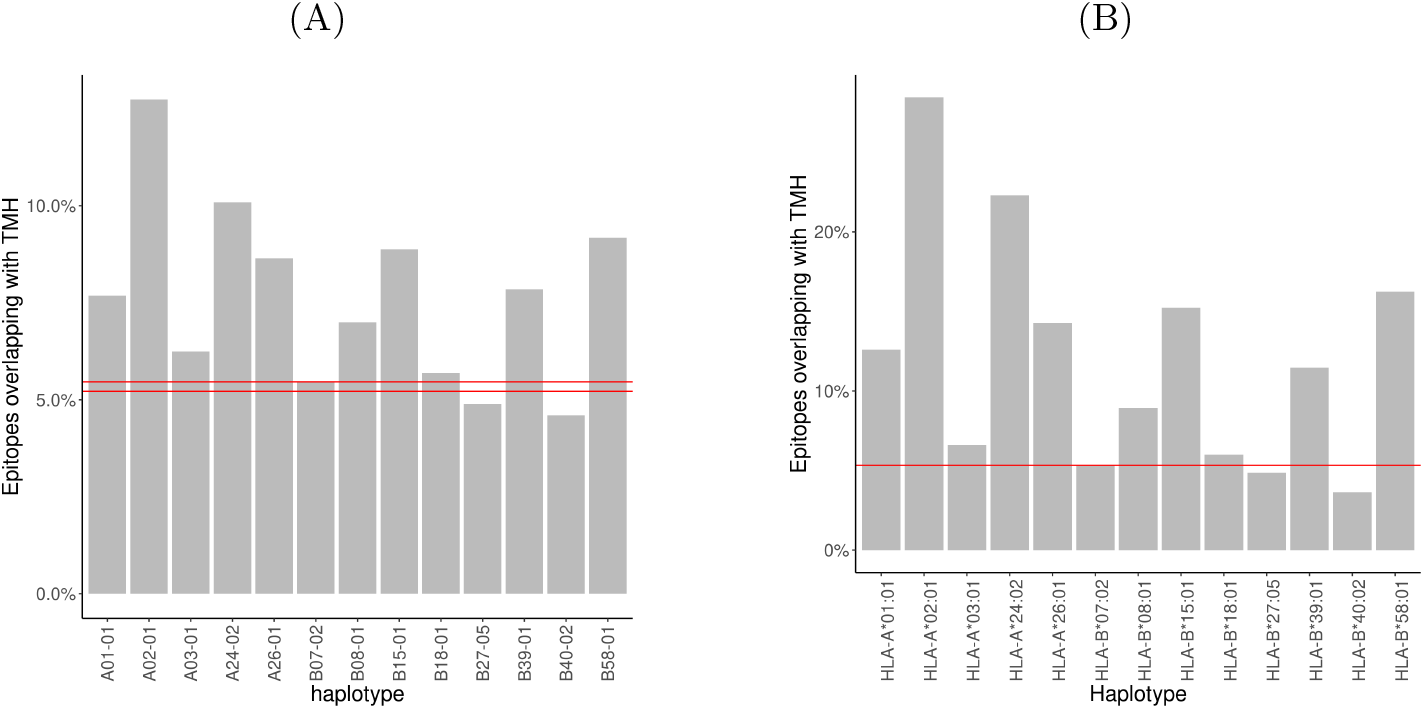
**(A)** Results for [7]. Red lines denotes the coincidence interval. **(B)** Results for this study. Red line denotes the percentage as expected by chance.

**Table 1:** Overview of all software used in this research.

| Goal | Tool | Reference |
| --- | --- | --- |
| Predict topology | TMHMM | [8] |
| Predict topology | PureseqTM | [13] |
| Predict epitopes MHC-I | **epitope-prediction** | [7] |
| Predict epitopes MHC-II | NetMHCIIpan | [43, 44] |
| Call TMHMM from R | tmhmm | [45] |
| Call PureseqTM from R | pureseqtmr | [46] |
| Call NetMHCIIpan from R | netmhc2pan | [47] |
| Combine all | bbbq | [48] |

### A.2 Prediction software used

For this research, we needed software to predict protein topology, as well as the MHC-I and MHC-II binding affinities of epitopes. We selected our software, by searching the scientific literature to identify the most recent free and open source (FOSS) prediction software. This was done by searching for papers that (1) cite older prediction software, and (2) present a novel method to make predictions. As a starting point, per type of prediction software, a review paper was used ([49] for protein topology, [50] for MHC-I binding affinities and [51] for MHC-II binding affinities).

There are multiple computational tools developed to predict which parts of a protein forms a TMH. In 2001, multiple of such prediction tools have been compared [49], of which TMHMM [8] turned out to be the most accurate, as is used in the previous study [7]. However, TMHMM has a restrictive software license and is nearly two decades old. Therefore, PureseqTM [13], was also used in this study, which has been more recently developed and has a free software license.

For MHC-I, there are multiple computational tools developed to predict epitopes. According to [50], at that time, NetMHCcons [52] gave the best predictions. We used the same tool as used in our earlier study, epitope-prediction [7],

Also for MHC-II, there are multiple computational tools developed to predict epitopes, such as using a trained neural network [51] or a Gibbs sampling approach [53]. According to [50], in 2011, from a set of multiple tools, NetMHCIIpan [43, 44] made the most accurate predictions. The most recent FOSS tool available now appears to be MHCnuggets [18], which can do both MHC-I and MHC-II predictions. As we already use epitope-prediction [7] for MHC-I predictions, we use MHCnuggets only for MHC-II predictions.

To retrieve the data from the NCBI databases the rentrez R package [54] was used that calls the NCBI website’s API. To provide for a stable user experience for all users, this API limits the user to 3 calls per second. Additionally, the API splits the result of a bigger query into multiple pages, each of which needs one API call. We wrote the sprentrez package [55] to provide for bigger queries of multiple (and delayed) API calls.

### A.3 Prediction software written

The R programming language is used for the complete experiment, including the analysis. The complete experiment is bundled in the ’bbbq’ R package, which is dependent on ’tmhmm’, ’pureseqtmr’, ’epitope-prediction’ and ’mhcnuggetsr’ as described below.

The R package ’tmhmm’ was developed to do the similar topology predictions as our earlier study (that used ’TMHMM’), yet in an automated way. ’TMHMM’ has a restrictive software license [8] and allows a user to download a pre-compiled executable after confirmation that he/she is in academia. The R package respects this restriction and allows the user to install and use TMHMM from within R, as done in this study. ’tmhmm’ has been submitted to and is accepted by the Comprehensive R Archive Network (CRAN).

To be able to call, from R, the TMH prediction software ’PureseqTM’ [13], which is written in C, the package ’pureseqtmr’ has been developed. ’pureseqtmr’ allows to install ’PureseqTM’ and use most of its features. ’pureseqtmr’ has been submitted to and is accepted by CRAN.

MHCnuggets is a free and open-source Python package to predict epitope affinity for many MHC-I and MHC-II variants [18]. The R package ’mhcnuggetsr’ allows one to install and use MHCnuggets from within R. Also ’mhcnuggetsr’ has been submitted to and is accepted by CRAN.

To reproduce the full experiment presented in this paper, the functions needed are bundled in the ’bbbq’ R package. This package is too specific to be submitted to CRAN.

**Table 2:** Percentage of spots and spots that overlap with a TMH

| target | mhc_class | n_spots | n_spots_tmh | f_tmh |
| --- | --- | --- | --- | --- |
| covid | 1 | 14207 | 1124 | 7.91 |
| covid | 2 | 14137 | 1245 | 8.81 |
| human | 1 | 11220940 | 598391 | 5.33 |
| human | 2 | 11118448 | 672273 | 6.05 |
| myco | 1 | 1299707 | 98613 | 7.59 |
| myco | 2 | 1279742 | 108419 | 8.47 |

### A.4 Prediction of percentage of epitopes overlapping with a TMH

Supplementary Table 2 shows an overview of the findings, where a target specifies the source of the proteome, where covid denotes SARS-CoV-2 and myco denotes *Mycobacterium tuberculosis*. mhc_class denotes the MHC class, n_spots the number of possible 9-mers (for MHC-I) or 14-mers (for MHC-II) possible. n_spots_tmh the number of epitopes that overlapped with a TMH that were binders. f_tmh the percentage of peptides that had at least 1 residue overlapping with a TMH.

### A.5 Minor methods

These are details that are removed from the ’Methods’ section.

PureseqTM does not predict the topology of proteins that have less than three amino acids. The TRDD1 (’T cell receptor delta diversity 1’) protein, however, is two amino acids long. The R package pureseqtmr, however, predicts that mono- and di-peptides are cytosolic.

### A.6 Minor discussion

These are details that are removed from the ’Discussion’ section.

In this experiment we predicted epitopes that overlap with TMHs from a human, bacterial and viral proteome, would these proteins be expressed in a human host. Bacteria, however have different cell membranes and cell walls, hence different structural requirements for a TMH. Both topology prediction tools were trained to recognize human TMHs, thus we cannot be sure that the transmembrane regions predicted in bacterial proteins are actually part of a TMH. For the purpose of this study, we assume the error in topology predictions to be unbiased way towards topology. In other words: that a bacterial TMH is incorrectly predicted to be absent just as often as it is incorrectly predicted to be present elsewhere.

Regarding the evolutionary conservation of TMHs using SNPs, again, it is estimated that approximately ten percent of SNPs is a false positive that result from the methods to determine a SNP. One example is that sequence variations are incorrectly detected due to highly similar duplicated sequences [56]. We assume that these duplications occur as often in TMHs as in regions around these, hence we expect this not to affect our results.

In our evolutionary experiment, we removed variations that were synonymous mutations (i.e. resulted in the same amino acid, from a different genetic code) from our analysis. There is evidence, however, that these synonymous mutations do have an effect and may even be evolutionary selected for [57]. As the possible effect of synonymous mutations is ignored by our topology prediction software, we do so as well.

### A.7 Elution studies

### A.8 IC50 values of binders per haplotype

Per target proteome (i.e. human, SARS-CoV-2, *M tuberculosis*), we collected all 9-mers (for MHC-I) and 14-mers (for MHC-II), after removing the selenoproteins and proteins that are shorter than the epitope length. From these epitopes, per MHC haplotype, we predicted the IC50 (in nM) using epitope-prediction (for MHC-I) and MHCnuggets (for MHC-II). Here, we show the IC50 value per haplotype that is used to determine if a peptide binds to the haplotype’s MHC for MHC-I (see supplementary Table 4) and MHC-II (see supplementary Table 5).

**Table 3:** Percentage of epitopes derived from a TMH found in the two elution studies, for the two different kind of topology prediction tools. The values between braces show the the number of epitopes that were predicted to overlapping with a TMH per all epitopes that could be uniquely mapped to the representative human reference proteome.

| MHC class | Tool | n |
| --- | --- | --- |
| I | PureseqTM | 1.38% (109/7897) |
| I | TMHMM | 1.43% (113/7897) |
| II | PureseqTM | 3.92% (498/12712) |
| II | TMHMM | 3.96% (504/12712) |

**Table 4:** IC50 values (in nM) per haplotype below which a peptide is considered a binder. percentage used: 2

| haplotype | covid | human | myco |
| --- | --- | --- | --- |
| HLA-A*01:01 | 1470.5912 | 2545.9537 | 2812.1714 |
| HLA-A*02:01 | 118.9596 | 218.7274 | 186.7565 |
| HLA-A*03:01 | 537.0144 | 804.7455 | 1544.1073 |
| HLA-A*24:02 | 984.8147 | 1590.0623 | 1971.8258 |
| HLA-A*26:01 | 1095.2591 | 1771.6924 | 1526.1101 |
| HLA-B*07:02 | 1215.7734 | 705.6514 | 435.5361 |
| HLA-B*08:01 | 886.5661 | 883.0951 | 1023.2213 |
| HLA-B*18:01 | 921.4157 | 1063.2215 | 1319.0445 |
| HLA-B*27:05 | 1186.0963 | 689.8815 | 475.6130 |
| HLA-B*39:01 | 437.3506 | 484.3843 | 399.3873 |
| HLA-B*40:02 | 585.6308 | 541.2392 | 600.1688 |
| HLA-B*58:01 | 435.4693 | 591.0526 | 538.9063 |
| HLA-B*15:01 | 281.9129 | 440.6541 | 482.8369 |

### A.9 Presentation of TMH-derived epitopes

Supplementary Table 6 shows the shorthand notation for the HLA haplotypes.

Supplementary Tables 7 and 8 show the exact number of binders, binders that overlap with TMHs and the percentage of binders that overlap with TMHs, as visualized by figure 1A.

**Table 5:**
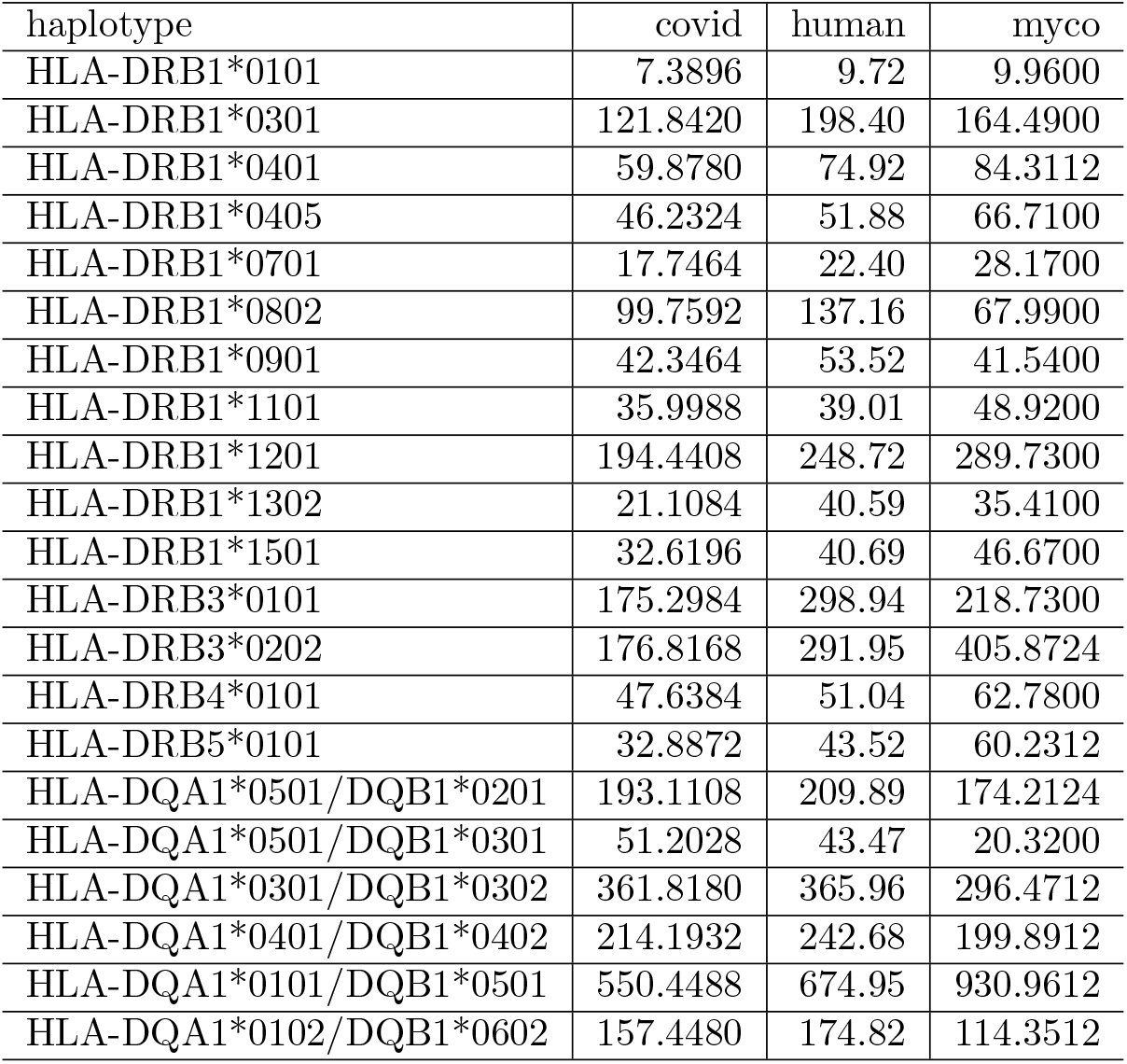
IC50 values (in nM) per haplotype below which a peptide is considered a binder. percentage used: 2

### A.10 Relative presentation of TMH-derived epitopes

To compare the over-presentation of TMH-derived epitopes between the different proteomes, we normalized this percentages in such a way that 1.0 is the percentage of TMH-derived epitopes that would be expected by chance. Figure 6 and 7 show these normalized values for the MHC-I and MHC-II haplotypes respectively.

**Figure 6:**
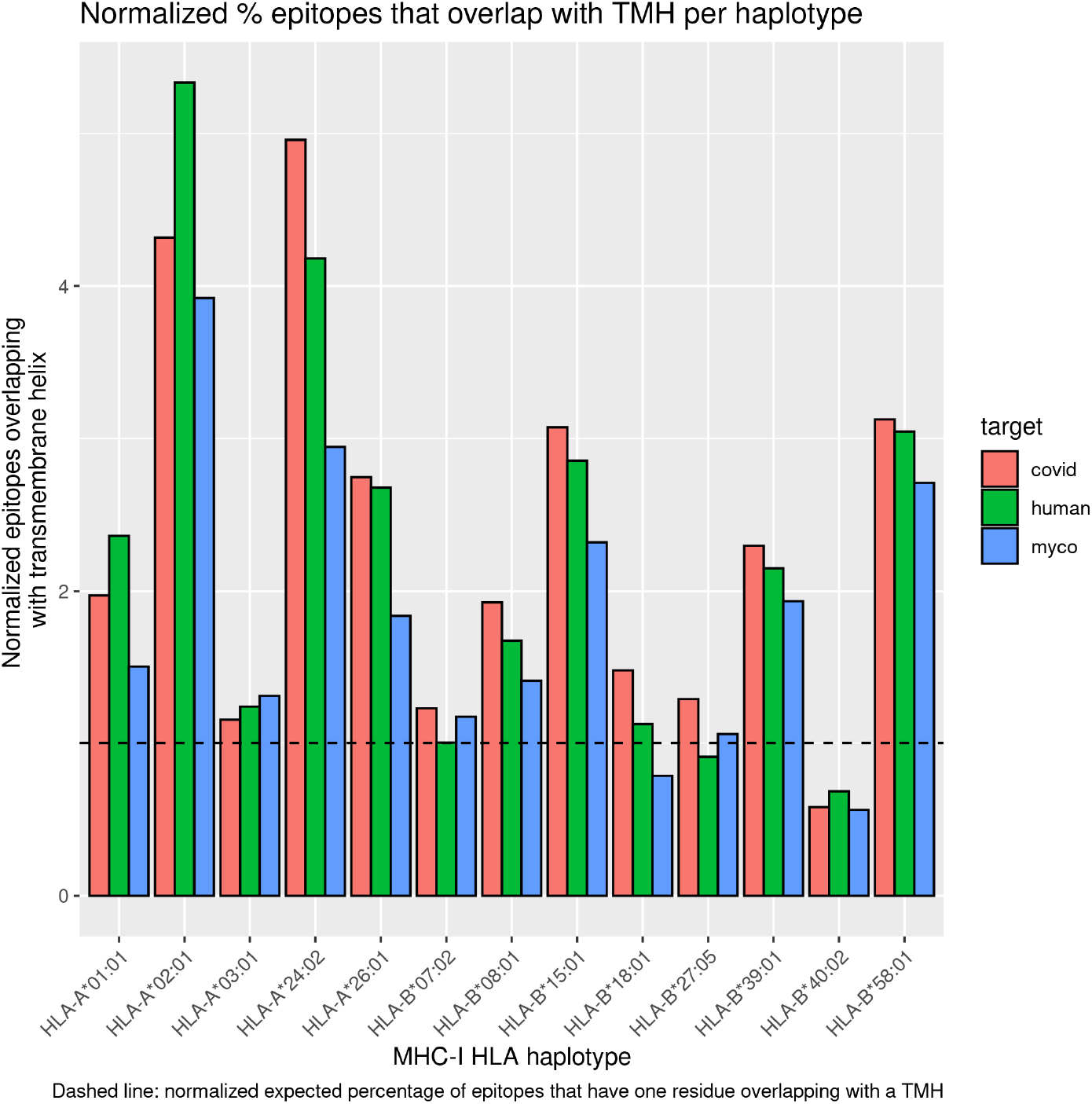
Normalized proportion of MHC-I epitopes overlapping with TMHs for human, viral and bacterial proteomes. Legend: covid = SARS-CoV-2, human = *Homo sapiens*, myco = *Mycobacterium tuberculosis*

**Figure 7:**
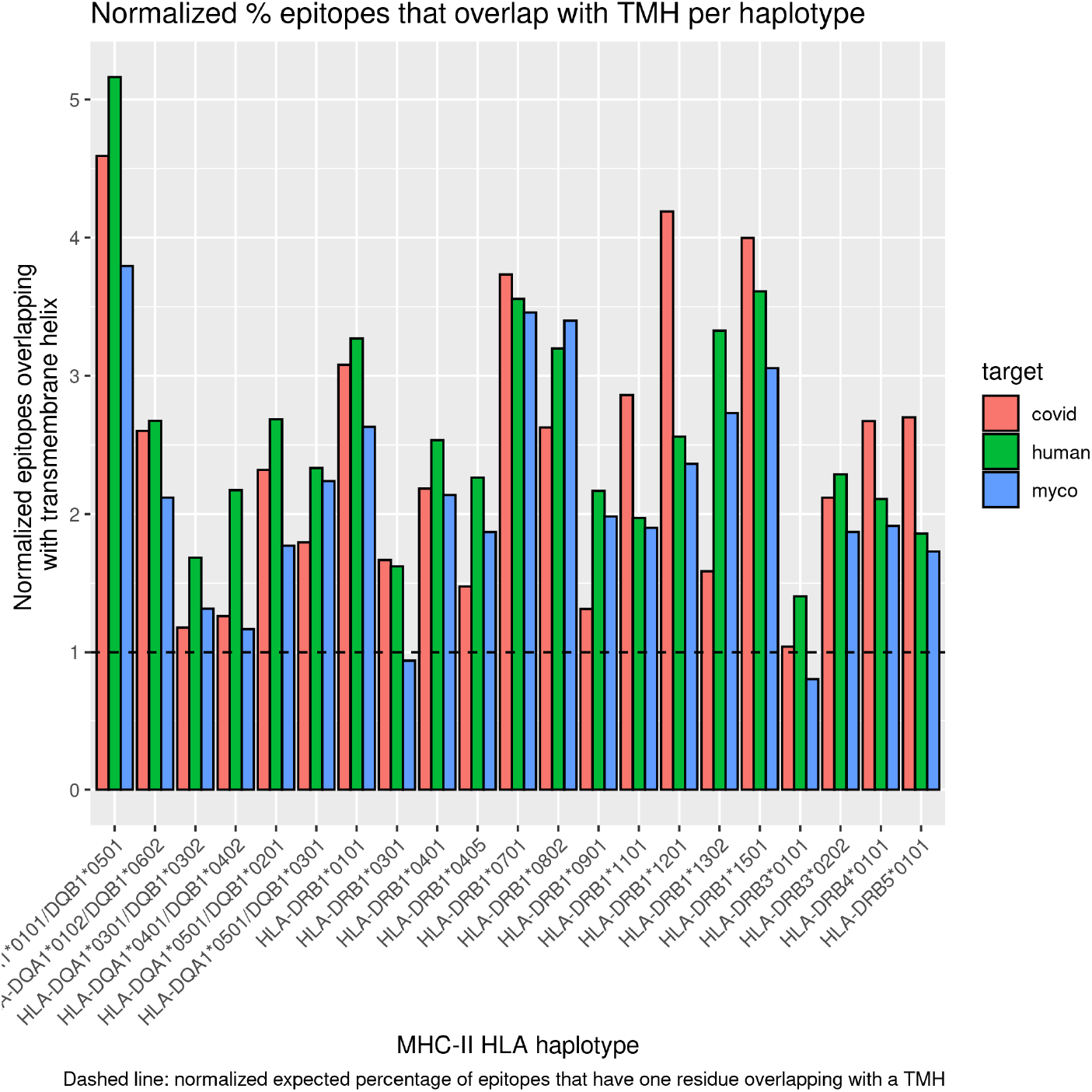
Normalized proportion of MHC-II epitopes overlapping with TMHs for human, viral and bacterial proteomes. Legend: covid = SARS-CoV-2, human = *Homo sapiens*, myco = *Mycobacterium tuberculosis*

To determine the additional over-presentation of TMH-derived epitopes in MHC-II (as compared to MHC-I), we normalized the data to enable a side-by-side comparison. The percentage of TMH-derived epitopes presented was normalized to the expected percentage of TMH-derived epitopes, where 1.0 denotes that the percentage of presented TMH-derived epitopes matches the values as expected by chance. The normalized values per haplotype are shown in figure 8. To compare the TMH-derived over-presentation per MHC class, we grouped the normalized values per haplotype, and plot the mean and standard error, as shown in figure 9.

**Figure 8:**
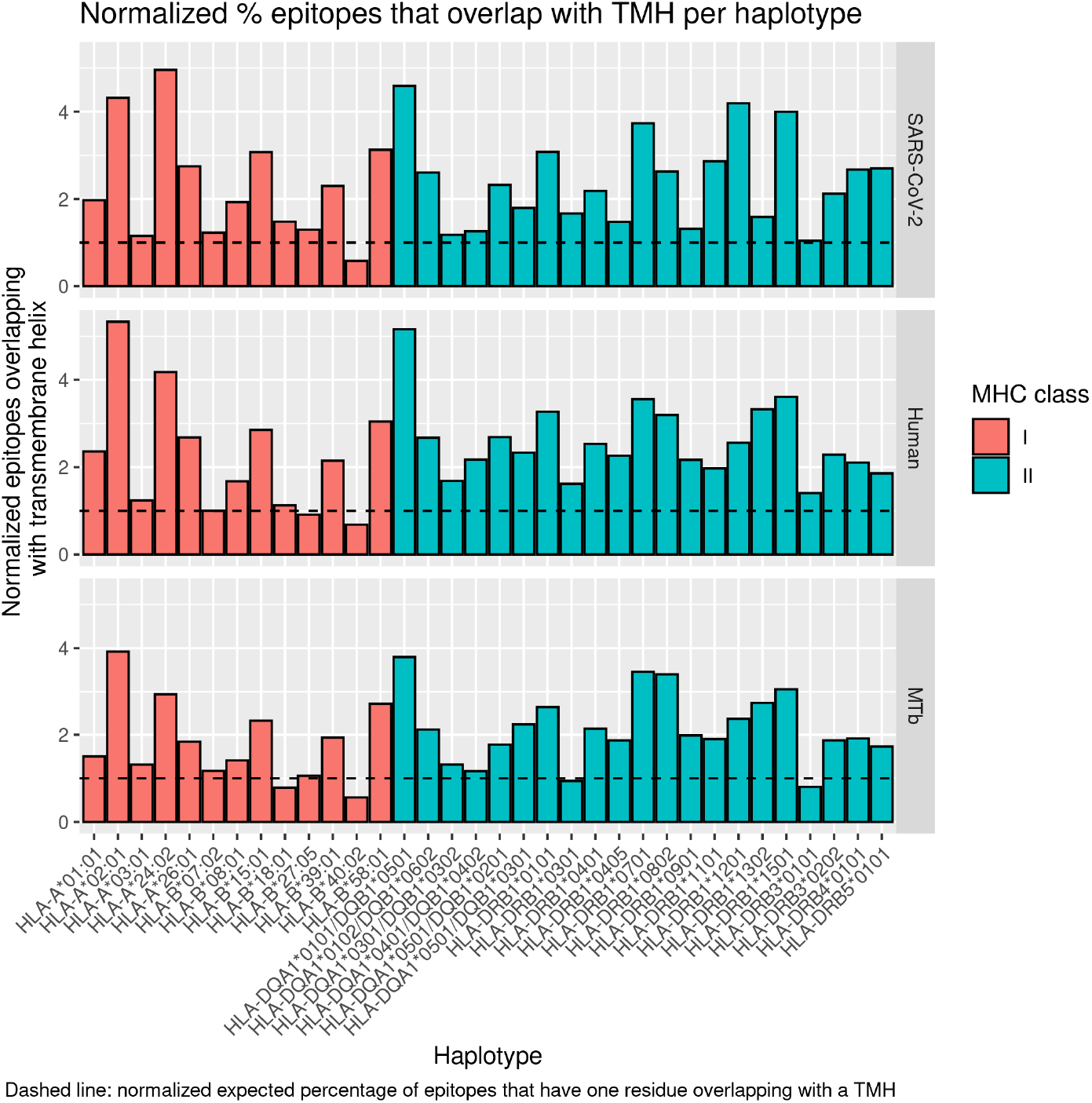
Normalized proportion of MHC-I and MHC-II epitopes overlapping with TMHs, for the different haplotypes and proteomes

**Figure 9:**
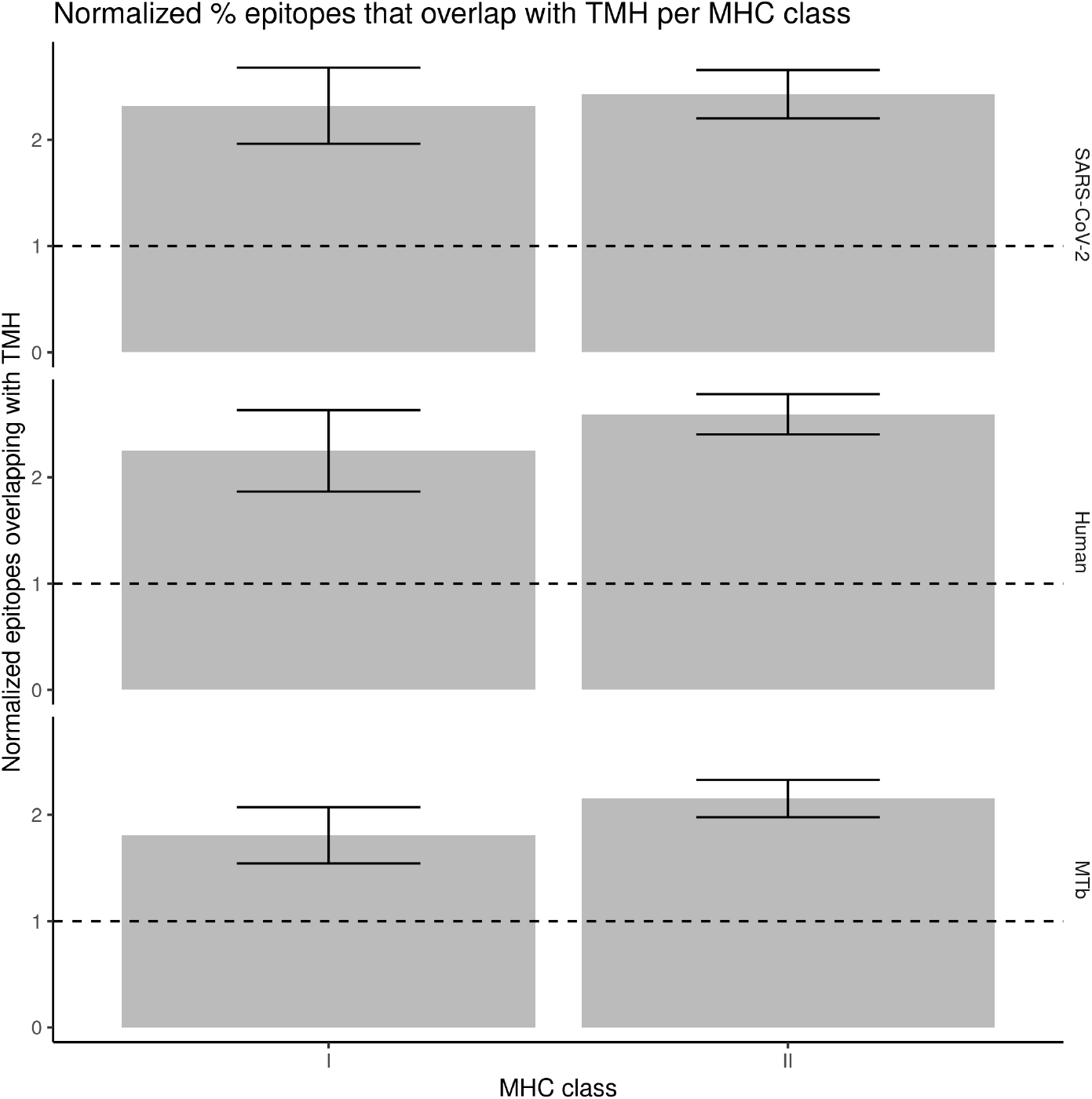
Normalized proportion of MHC-I and MHC-II epitopes overlapping with TMHs, for the different MHC classes and proteomes. Error bars denote the standard error.

### A.11 Evolutionary conservation

See supplementary Tables 9 and 10 for an overview of all amounts. In supplementary Table 9 there are multiple instances where the amounts are expected to add up, yet don’t, as one SNP can work on multiple isoforms. For example, there are 9,621 unique SNPs found in all proteins, of which 4,219 around found in MAPs and 6,026 in TMPs. Apparently, 624 SNPs work on a set of isoforms that contains both MAPs and TMPs.

Figure 10 shows the distribution of the number of SNPs per gene name, at the date we started the experiment, at December 14th 2020.

**Figure 10:**
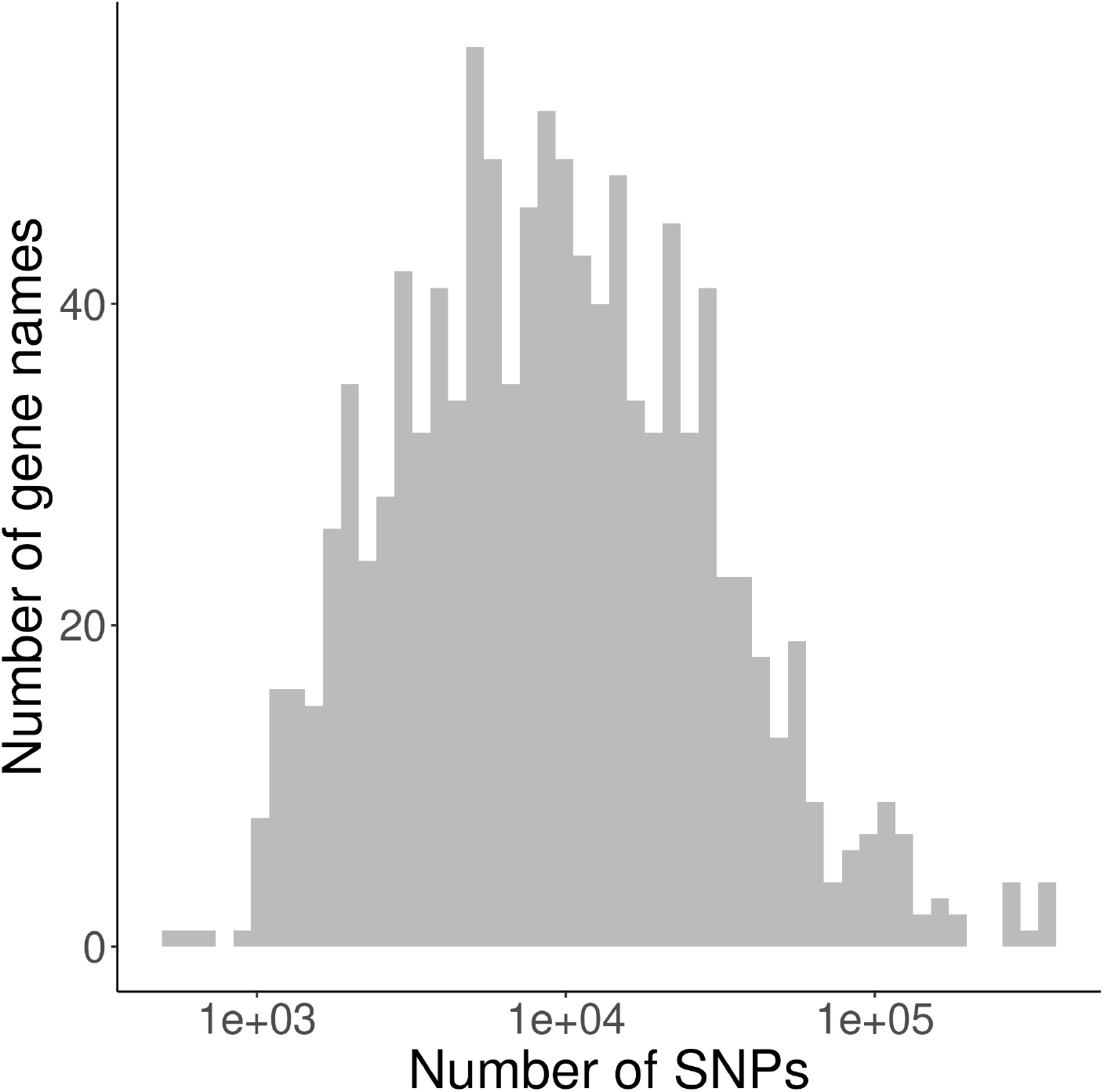
Distribution of the number of SNPs per gene name in the NCBI database.

**Figure 11:**
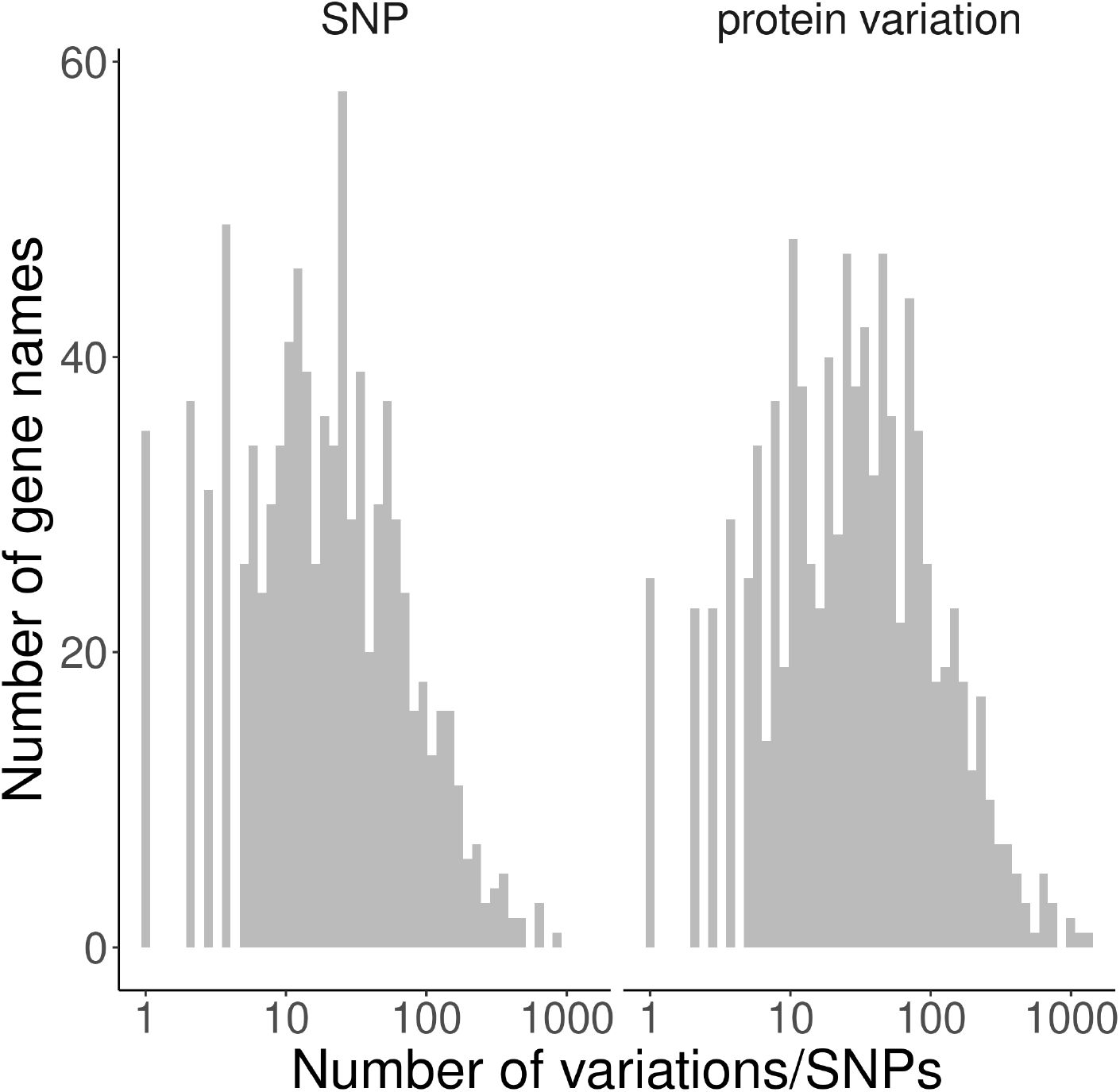
Distribution of the number of protein variations and SNPs per gene name processed.

**Figure 12:**
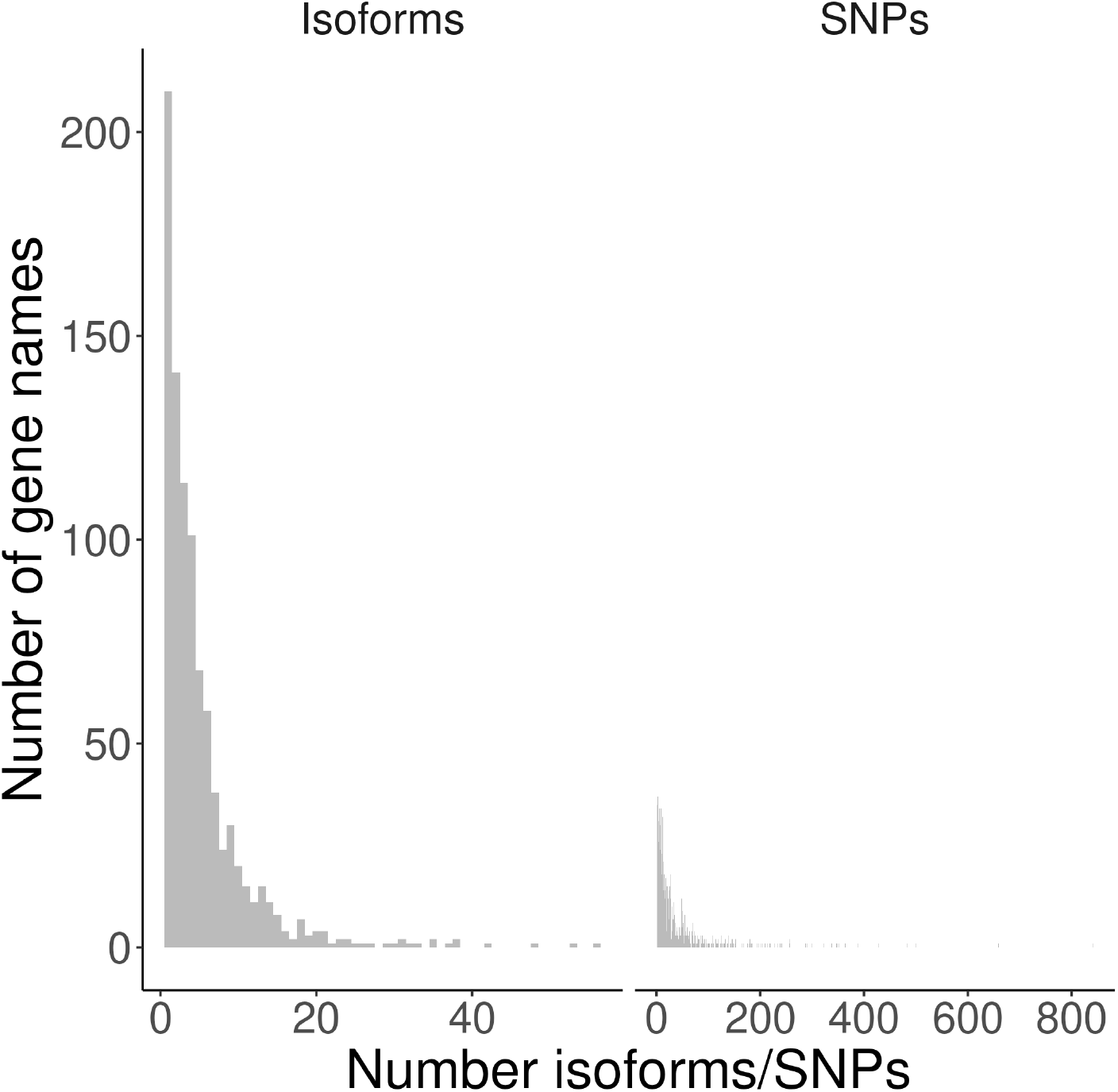
Histogram of the number of proteins found per gene name. Most often, a gene name is associated with one proteins.

**Figure 13:**
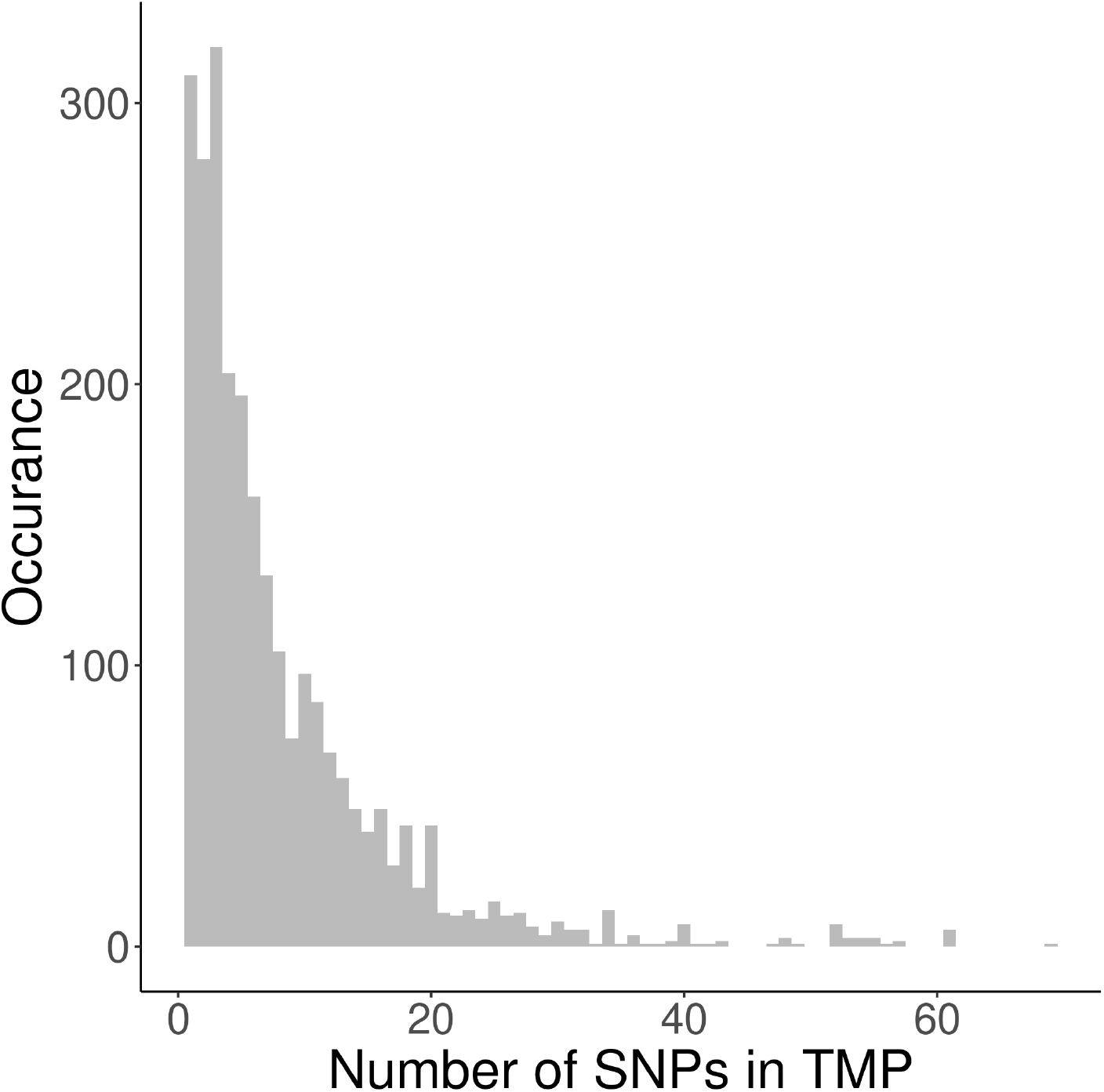
Histogram of the number of SNPs per trans-membrane protein. Dashed vertical line: average number of SNPs per TMP

**Figure 14:**
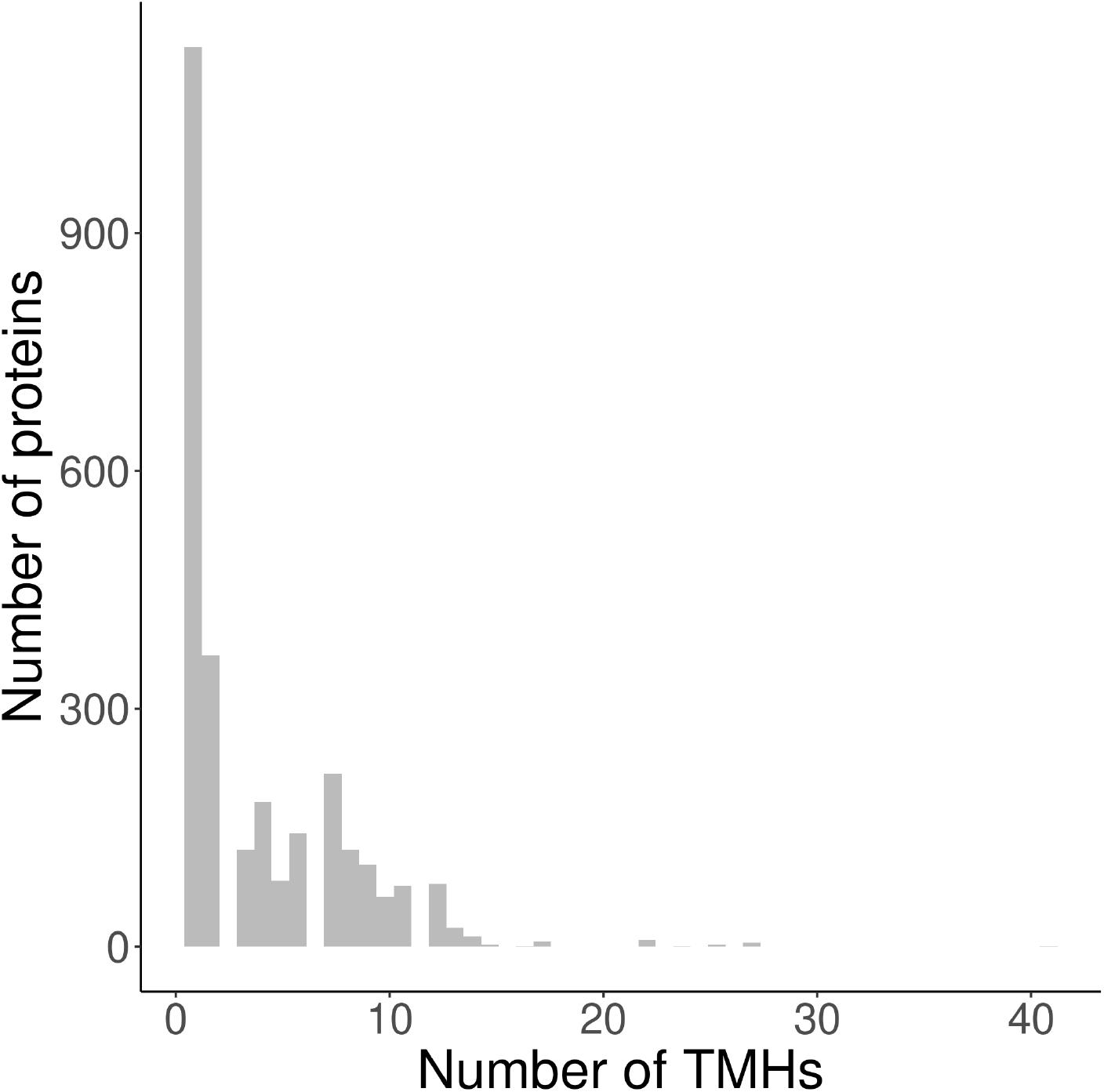
Histogram of the number of TMHs predicted per protein, for the trans-membrane proteins used.

To verify if SNPs were sampled uniformly over proteins, we show the distribution of the relative position in figure 15. We find no clear evidence of a bias.

**Figure 15:**
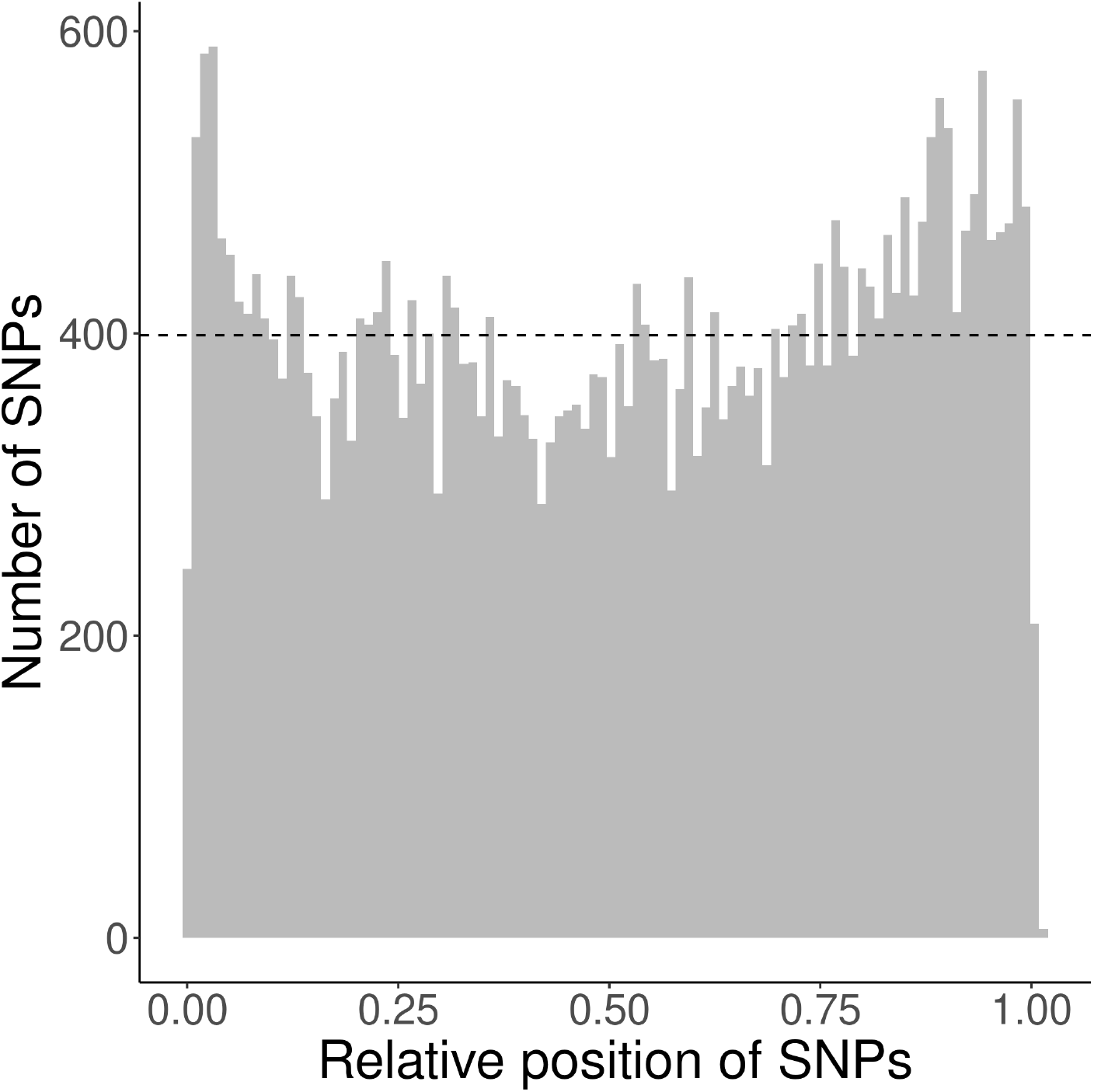
Distribution of the relative position of the SNPs used, where a relative position of zero denotes the first amino acid at the N-terminus, where a relative position of one indicates the last residue at the C-terminus.

Supplementary Table 11 shows the statistics for all SNPs, where supplementary Tables 12 and 13 show the statistics for only single-spanners and multi-spanners respectively.

**Table 6:**
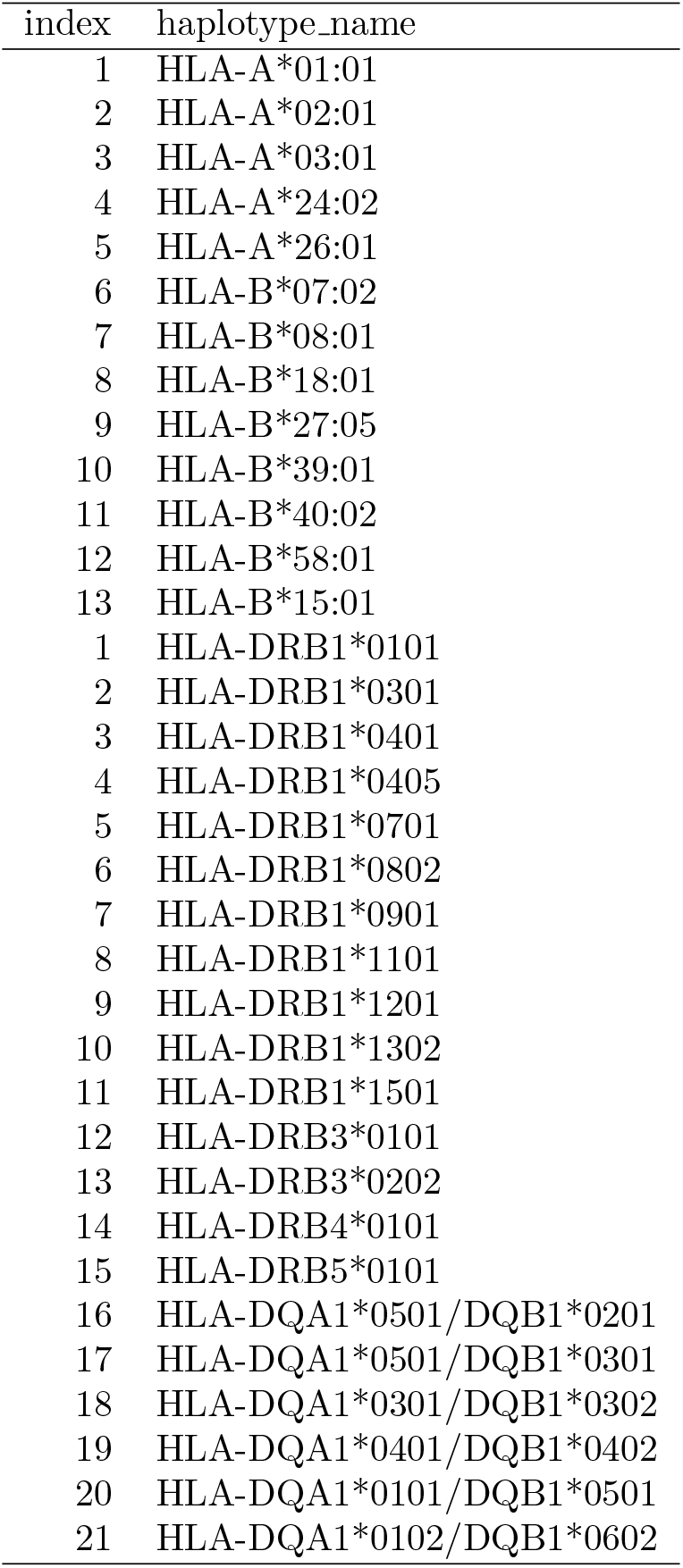
Abbreviations of the haplotype names

**Table 7:**
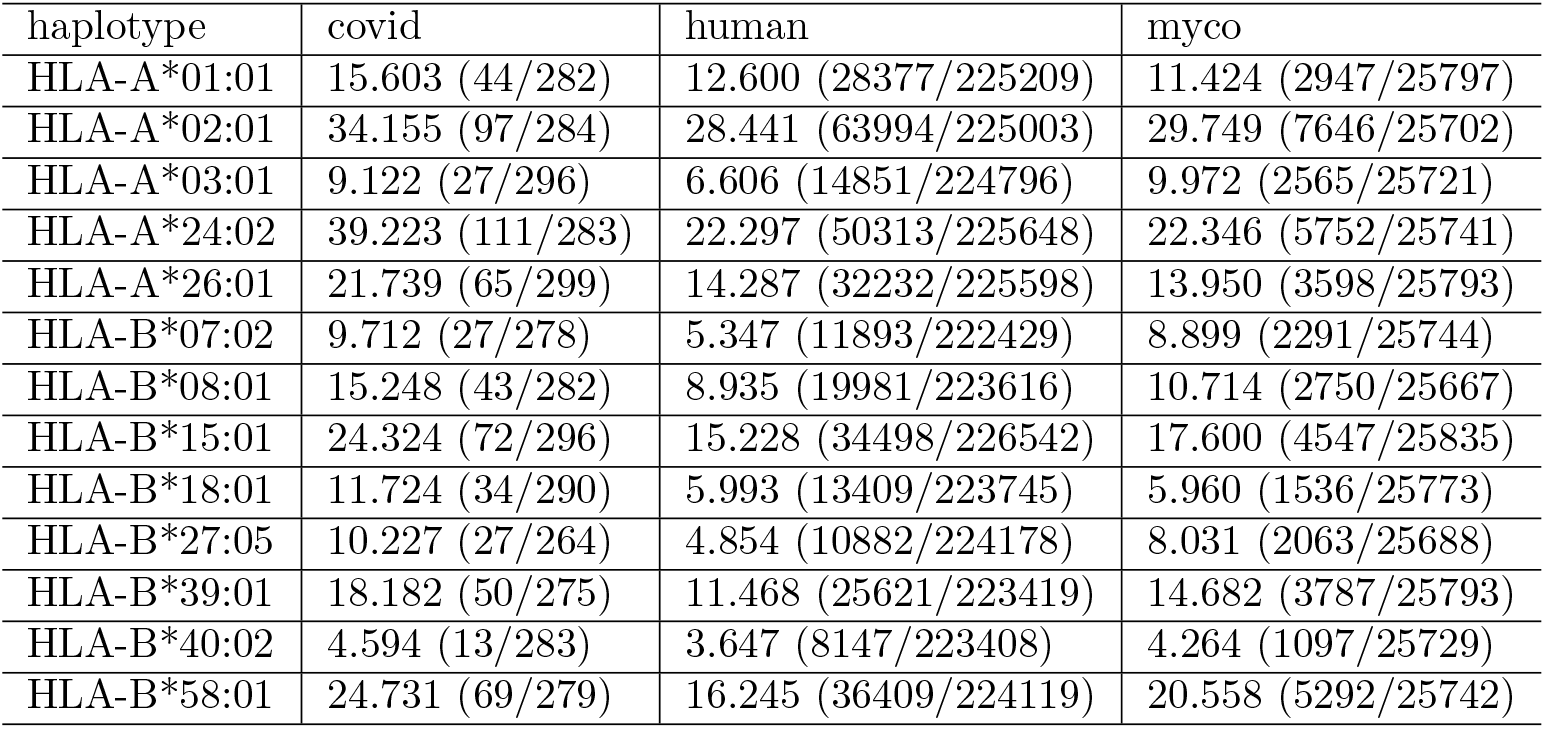
Percentage of MHC-I 9-mers overlapping with TMH. Values in brackets show the number of binders that have at least one residue overlapping with a TMH (first value)as well as the number of binders (second value). percentage used: 2

| haplotype | covid | human | myco |
| --- | --- | --- | --- |
| HLA-A*01:01 | 15.603 (44/282) | 12.600 (28377/225209) | 11.424 (2947/25797) |
| HLA-A*02:01 | 34.155 (97/284) | 28.441 (63994/225003) | 29.749 (7646/25702) |
| HLA-A*03:01 | 9.122 (27/296) | 6.606 (14851/224796) | 9.972 (2565/25721) |
| HLA-A*24:02 | 39.223 (111/283) | 22.297 (50313/225648) | 22.346 (5752/25741) |
| HLA-A*26:01 | 21.739 (65/299) | 14.287 (32232/225598) | 13.950 (3598/25793) |
| HLA-B*07:02 | 9.712 (27/278) | 5.347 (11893/222429) | 8.899 (2291/25744) |
| HLA-B*08:01 | 15.248 (43/282) | 8.935 (19981/223616) | 10.714 (2750/25667) |
| HLA-B*15:01 | 24.324 (72/296) | 15.228 (34498/226542) | 17.600 (4547/25835) |
| HLA-B*18:01 | 11.724 (34/290) | 5.993 (13409/223745) | 5.960 (1536/25773) |
| HLA-B*27:05 | 10.227 (27/264) | 4.854 (10882/224178) | 8.031 (2063/25688) |
| HLA-B*39:01 | 18.182 (50/275) | 11.468 (25621/223419) | 14.682 (3787/25793) |
| HLA-B*40:02 | 4.594 (13/283) | 3.647 (8147/223408) | 4.264 (1097/25729) |
| HLA-B*58:01 | 24.731 (69/279) | 16.245 (36409/224119) | 20.558 (5292/25742) |

**Table 8:** Percentage of MHC-II 14-mers overlapping with TMH. Values in brackets show the number of binders that have at least one residue overlapping with a TMH (first value)as well as the number of binders (second value). percentage used: 2

| haplotype | covid | human | myco |
| --- | --- | --- | --- |
| HLA-DQA1*0101/DQB1*0501 | 40.433 (112/277) | 31.214 (69752/223464) | 32.158 (8187/25459) |
| HLA-DQA1*0102/DQB1*0602 | 22.910 (74/323) | 16.167 (35753/221147) | 17.950 (4608/25671) |
| HLA-DQA1*0301/DQB1*0302 | 10.381 (30/289) | 10.179 (22623/222248) | 11.144 (2842/25502) |
| HLA-DQA1*0401/DQB1*0402 | 11.111 (32/288) | 13.135 (29319/223219) | 9.890 (2524/25522) |
| HLA-DQA1*0501/DQB1*0201 | 20.430 (57/279) | 16.240 (36186/222820) | 14.999 (3823/25489) |
| HLA-DQA1*0501/DQB1*0301 | 15.808 (46/291) | 14.106 (31046/220089) | 18.969 (4878/25715) |
| HLA-DRB1*0101 | 27.119 (80/295) | 19.774 (43968/222349) | 22.293 (5692/25533) |
| HLA-DRB1*0301 | 14.676 (43/293) | 9.801 (21831/222752) | 7.956 (2025/25451) |
| HLA-DRB1*0401 | 19.231 (55/286) | 15.325 (34011/221930) | 18.113 (4641/25623) |
| HLA-DRB1*0405 | 12.996 (36/277) | 13.684 (30380/222012) | 15.837 (4036/25484) |
| HLA-DRB1*0701 | 32.877 (96/292) | 21.512 (47856/222465) | 29.304 (7471/25495) |
| HLA-DRB1*0802 | 23.132 (65/281) | 19.339 (42859/221623) | 28.805 (7358/25544) |
| HLA-DRB1*0901 | 11.565 (34/294) | 13.111 (29043/221520) | 16.798 (4301/25605) |
| HLA-DRB1*1101 | 25.197 (64/254) | 11.924 (26582/222928) | 16.103 (4101/25467) |
| HLA-DRB1*1201 | 36.897 (107/290) | 15.482 (34596/223464) | 20.018 (5098/25467) |
| HLA-DRB1*1302 | 13.962 (37/265) | 20.121 (44798/222646) | 23.141 (5935/25647) |
| HLA-DRB1*1501 | 35.206 (94/267) | 21.836 (48671/222893) | 25.891 (6584/25430) |
| HLA-DRB3*0101 | 9.158 (25/273) | 8.496 (18884/222274) | 6.819 (1740/25517) |
| HLA-DRB3*0202 | 18.657 (50/268) | 13.832 (30687/221859) | 15.843 (4059/25620) |
| HLA-DRB4*0101 | 23.529 (68/289) | 12.749 (28376/222568) | 16.221 (4131/25467) |
| HLA-DRB5*0101 | 23.776 (68/286) | 11.235 (24993/222464) | 14.648 (3732/25478) |

**Table 9:**
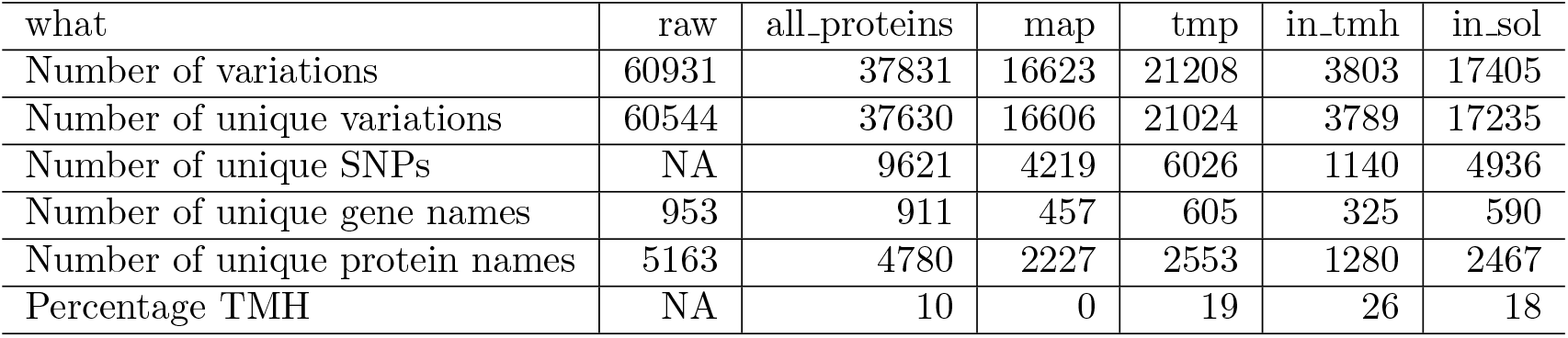
Amounts. raw = all variations, including DNA variations. all_proteins = all proteins. map = membrane associated protein. tmp = transmembrane protein. in_tmh = in transmembrane helix of TMP. in_sol = in soluble region of TMP.

**Table 10:** Amounts. single_in_tmh = in transmembrane helix of single-spanner. single_in_sol = in soluble region of single-spanner. multi_in_tmh = in transmembrane helix of multi-spanner. multi_in_sol = in soluble region of multi-spanner.

| what | single_in_tmh | single_in_sol | multi_in_tmh | multi_in_sol |
| --- | --- | --- | --- | --- |
| Number of variations | 452 | 7734 | 3351 | 9671 |
| Number of unique variations | 451 | 7733 | 3338 | 9502 |
| Number of unique SNPs | 160 | 2393 | 994 | 2762 |
| Number of unique gene names | 96 | 282 | 243 | 344 |
| Number of unique protein names | 304 | 1032 | 976 | 1435 |
| Percentage TMH | 11 | 5 | 35 | 26 |

**Table 11:** Statistics for all TMPs. p = p value. n = number of SNPs. n_success = number of SNPs found in TMHs (dashed blue line). E(n_success) = expected number of SNPs to be found in TMHs (dashed red line).

| parameter | value |
| --- | --- |
| p | 6.820823e-11 |
| n | 21208 |
| n_success | 3803 |
| E(n_success) | 4140.56 |

**Table 12:** Statistics for the single-spanners. p = p value. n = number of SNPs in single-spanners. n_success = number of SNPs found in TMHs of single-spanners (dashed blue line). E(n_success) = expected number of SNPs to be found in TMHs of single-spanners (dashed red line).

| parameter | value |
| --- | --- |
| p | 0.3189532 |
| n | 8186 |
| n_success | 452 |
| E(n_success) | 462.1535 |

**Table 13:**
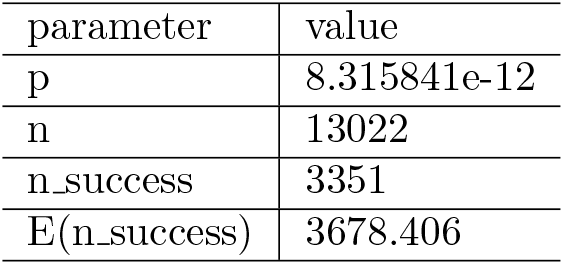
Statistics for the multi-spanners. p = p value. n = number of SNPs in multi-spanners. n_success = number of SNPs found in TMHs of multi-spanners (dashed blue line). E(n_success) = expected number of SNPs to be found in TMHs of multi-spanners (dashed red line).

## References

[1] Ole Lund, Morten Nielsen, Can Kesmir, Anders Gorm Petersen, Claus Lundegaard, Peder Worning, Christina Sylvester-Hvid, Kasper Lamberth, Gustav Røder, Sune Justesen, et al. Definition of supertypes for HLA molecules using clustering of specificity matrices. Immunogenetics, 55(12): 797–810, 2004.

[2] Steven GE Marsh, ED Albert, WF Bodmer, RE Bontrop, B Dupont, HA Erlich, M Fernández-Viña, DE Geraghty, R Holdsworth, CK Hurley, et al. Nomenclature for factors of the HLA system, 2010. Tissue antigens, 75(4):291, 2010.

[3] Simone Sommer. The importance of immune gene variability (MHC) in evolutionary ecology and conservation. Frontiers in zoology, 2(1):1–18, 2005.

[4] Mette Voldby Larsen, Alina Lelic, Robin Parsons, Morten Nielsen, Ilka Hoof, Kasper Lamberth, Mark B Loeb, Søren Buus, Jonathan Bramson, and Ole Lund. Identification of CD8+ T cell epitopes in the West Nile virus polyprotein by reverse-immunology using NetCTL. PloS one, 5(9), 2010.

[5] Ingrid MM Schellens, Can Kesmir, Frank Miedema, Debbie van Baarle, and José AM Borghans. An unanticipated lack of consensus cytotoxic T lymphocyte epitopes in HIV-1 databases: the contribution of prediction programs. Aids, 22(1):33–37, 2008.

[6] Sheila T Tang, Krista E van Meijgaarden, Nadia Caccamo, Giuliana Guggino, Michél R Klein, Pascale van Weeren, Fatima Kazi, Anette Stryhn, Alexander Zaigler, Ugur Sahin, et al. Genome-based in silico identification of new Mycobacterium tuberculosis antigens activating polyfunctional CD8+ T cells in human tuberculosis. The Journal of Immunology, 186(2): 1068–1080, 2011.

[7] Frans Bianchi, Johannes Textor, and Geert van den Bogaart. Transmembrane helices are an overlooked source of Major Histocompatibility Complex Class I epitopes. Frontiers in immunology, 8:1118, 2017.

[8] Anders Krogh, Björn Larsson, Gunnar Von Heijne, and Erik LL Sonnhammer. Predicting transmembrane protein topology with a hidden Markov model: application to complete genomes. Journal of molecular biology, 305 (3):567–580, 2001.

[9] Lukas Käll, Anders Krogh, and Erik LL Sonnhammer. A combined trans-membrane topology and signal peptide prediction method. Journal of molecular biology, 338(5):1027–1036, 2004.

[10] Masafumi Arai, Hironori Mitsuke, Masami Ikeda, Jun-Xiong Xia, Takashi Kikuchi, Masanobu Satake, and Toshio Shimizu. ConPred II: a consensus prediction method for obtaining transmembrane topology models with high reliability. Nucleic acids research, 32(suppl 2):W390–W393, 2004.

[11] David T Jones. Improving the accuracy of transmembrane protein topology prediction using evolutionary information. Bioinformatics, 23(5):538–544, 2007.

[12] Martin Klammer, David N Messina, Thomas Schmitt, and Erik LL Sonnhammer. MetaTM-a consensus method for transmembrane protein topology prediction. BMC bioinformatics, 10(1):314, 2009.

[13] Qing Wang, Chongming Ni, Zhen Li, Xiufeng Li, Renmin Han, Feng Zhao, Jinbo Xu, Xin Gao, and Sheng Wang. PureseqTM: efficient and accurate prediction of transmembrane topology from amino acid sequence only. bioRxiv, page 627307, 2019.

[14] Mamoun Ahram, Zoi I Litou, Ruihua Fang, and Ghaith Al-Tawallbeh. Estimation of membrane proteins in the human proteome. In silico biology, 6(5):379–386, 2006.

[15] Tara Hessa, Nadja M Meindl-Beinker, Andreas Bernsel, Hyun Kim, Yoko Sato, Mirjam Lerch-Bader, IngMarie Nilsson, Stephen H White, and Gunnar Von Heijne. Molecular code for transmembrane-helix recognition by the sec61 translocon. Nature, 450(7172):1026–1030, 2007.

[16] DT Jones, WR Taylor, and JM Thornton. A model recognition approach to the prediction of all-helical membrane protein structure and topology. Biochemistry, 33(10):3038–3049, 1994.

[17] Elin Bergseng, Siri Dørum, Magnus Ø Arntzen, Morten Nielsen, Ståle Nygård, Søren Buus, Gustavo A de Souza, and Ludvig M Sollid. Different binding motifs of the celiac disease-associated hla molecules DQ2.5, DQ2.2, and DQ7.5 revealed by relative quantitative proteomics of endogenous peptide repertoires. Immunogenetics, 67(2):73–84, 2015.

[18] Xiaoshan M Shao, Rohit Bhattacharya, Justin Huang, IK Ashok Sivakumar, Collin Tokheim, Lily Zheng, Dylan Hirsch, Benjamin Kaminow, Ashton Omdahl, Maria Bonsack, et al. High-throughput prediction of MHC class I and II neoantigens with MHCnuggets. Cancer Immunology Research, 8(3):396–408, 2020.

[19] Jason Greenbaum, John Sidney, Jolan Chung, Christian Brander, Bjoern Peters, and Alessandro Sette. Functional classification of class II human leukocyte antigen (HLA) molecules reveals seven different supertypes and a surprising degree of repertoire sharing across supertypes. Immunogenetics, 63(6):325–335, 2011.

[20] Ingrid MM Schellens, Ilka Hoof, Hugo D Meiring, Sanne NM Spijkers, Martien CM Poelen, Kees van der Poel, Ana I Costa, Cecile ACM van Els, Debbie van Baarle, Can Kesmir, et al. Comprehensive analysis of the naturally processed peptide repertoire: differences between HLA-A and B in the immunopeptidome. PloS one, 10(9):e0136417, 2015.

[21] Stephen T Sherry, M-H Ward, M Kholodov, J Baker, Lon Phan, Elizabeth M Smigielski, and Karl Sirotkin. dbSNP: the ncbi database of genetic variation. Nucleic acids research, 29(1):308–311, 2001.

[22] Garth R Brown, Vichet Hem, Kenneth S Katz, Michael Ovetsky, Craig Wallin, Olga Ermolaeva, Igor Tolstoy, Tatiana Tatusova, Kim D Pruitt, Donna R Maglott, et al. Gene: a gene-centered information resource at NCBI. Nucleic acids research, 43(D1):D36–D42, 2015.

[23] Eric W Sayers, Tanya Barrett, Dennis A Benson, Evan Bolton, Stephen H Bryant, Kathi Canese, Vyacheslav Chetvernin, Deanna M Church, Michael DiCuccio, Scott Federhen, et al. Database resources of the national center for biotechnology information. Nucleic acids research, 39(suppl 1):D38–D51, 2010.

[24] Kenneth L Rock, Eric Reits, and Jacques Neefjes. Present yourself! by mhc class i and mhc class ii molecules. Trends in immunology, 37(11):724–737, 2016.

[25] Andreas Blees, Dovile Januliene, Tommy Hofmann, Nicole Koller, Carla Schmidt, Simon Trowitzsch, Arne Moeller, and Robert Tampé. Structure of the human mhc-i peptide-loading complex. Nature, 551(7681):525–528, 2017.

[26] G Michael Preston and Jeffrey L Brodsky. The evolving role of ubiquitin modification in endoplasmic reticulum-associated degradation. Biochemical Journal, 474(4):445–469, 2017.

[27] Birgit Meusser, Christian Hirsch, Ernst Jarosch, and Thomas Sommer. Erad: the long road to destruction. Nature cell biology, 7(8):766–772, 2005.

[28] Laurence Bougnères, Julie Helft, Sangeeta Tiwari, Pablo Vargas, Benny Hung-Junn Chang, Lawrence Chan, Laura Campisi, Gregoire Lauvau, Stephanie Hugues, Pradeep Kumar, et al. A role for lipid bodies in the cross-presentation of phagocytosed antigens by mhc class i in dendritic cells. Immunity, 31(2):232–244, 2009.

[29] Toyoshi Fujimoto and Yuki Ohsaki. The proteasomal and autophagic pathways converge on lipid droplets. Autophagy, 2(4):299–301, 2006.

[30] Clàudia C Oliveira and Thorbald van Hall. Alternative antigen processing for mhc class i: multiple roads lead to rome. Frontiers in immunology, 6: 298, 2015.

[31] Georg Lautscham, Sabine Mayrhofer, Graham Taylor, Tracey Haigh, Alison Leese, Alan Rickinson, and Neil Blake. Processing of a multiple membrane spanning epstein-barr virus protein for cd8+ t cell recognition reveals a proteasome-dependent, transporter associated with antigen processing–independent pathway. The Journal of experimental medicine, 194(8):1053–1068, 2001.

[32] Jean Gruenberg. Life in the lumen: the multivesicular endosome. Traffic, 21(1):76–93, 2020.

[33] Monique Kleijmeer, Georg Ramm, Danita Schuurhuis, Janice Griffith, Maria Rescigno, Paola Ricciardi-Castagnoli, Alexander Y Rudensky, Ferry Ossendorp, Cornelis JM Melief, Willem Stoorvogel, et al. Reorganization of multivesicular bodies regulates mhc class ii antigen presentation by dendritic cells. The Journal of cell biology, 155(1):53–64, 2001.

[34] Peter J Peters, Jacques J Neefjes, Viola Oorschot, Hidde L Ploegh, and Hans J Geuze. Segregation of mhc class ii molecules from mhc class i molecules in the golgi complex for transport to lysosomal compartments. Nature, 349(6311):669–676, 1991.

[35] Wilbert Zwart, Alexander Griekspoor, Coenraad Kuijl, Marije Marsman, Jacco van Rheenen, Hans Janssen, Jero Calafat, Marieke van Ham, Lennert Janssen, Marcel van Lith, et al. Spatial separation of hla-dm/hla-dr interactions within miic and phagosome-induced immune escape. Immunity, 22 (2):221–233, 2005.

[36] Peter Sander, Katja Becker, and Michael Dal Molin. Lipase processing of complex lipid antigens. Cell chemical biology, 23(9):1044–1046, 2016.

[37] Martine Gilleron, Marco Lepore, Emilie Layre, Diane Cala-De Paepe, Naila Mebarek, James A Shayman, Stèphane Canaan, Lucia Mori, Frèdèric Carrière, Germain Puzo, et al. Lysosomal lipases plrp2 and lpla2 process mycobacterial multi-acylated lipids and generate t cell stimulatory antigens. Cell chemical biology, 23(9):1147–1156, 2016.

[38] Ilse Dingjan, Daniëlle RJ Verboogen, Laurent M Paardekooper, Natalia H Revelo, Simone P Sittig, Linda J Visser, Gabriele Fischer Von Mollard, Stefanie SV Henriet, Carl G Figdor, Martin Ter Beest, et al. Lipid peroxidation causes endosomal antigen release for cross-presentation. Scientific reports, 6(1):1–12, 2016.

[39] Lauro Velazquez-Salinas, Selene Zarate, Samantha Eberl, Douglas P Gladue, Isabel Novella, and Manuel V Borca. Positive selection of ORF3a and ORF8 genes drives the evolution of SARS-CoV-2 during the 2020 COVID-19 pandemic. bioRxiv, 2020.

[40] Alvin X Han, Sebastian Maurer-Stroh, and Colin A Russell. Individual immune selection pressure has limited impact on seasonal influenza virus evolution. Nature ecology & evolution, 3(2):302–311, 2019.

[41] Timothy J Stevens and Isaiah T Arkin. Substitution rates in *α*-helical transmembrane proteins. Protein Science, 10(12):2507–2517, 2001.

[42] Amit Oberai, Nathan H Joh, Frank K Pettit, and James U Bowie. Structural imperatives impose diverse evolutionary constraints on helical membrane proteins. Proceedings of the National Academy of Sciences, 106(42): 17747–17750, 2009.

[43] Morten Nielsen, Claus Lundegaard, Thomas Blicher, Bjoern Peters, Alessandro Sette, Sune Justesen, Søren Buus, and Ole Lund. Quantitative predictions of peptide binding to any HLA-DR molecule of known sequence: NetMHCIIpan. PLoS computational biology, 4(7), 2008.

[44] Edita Karosiene, Michael Rasmussen, Thomas Blicher, Ole Lund, Søren Buus, and Morten Nielsen. NetMHCIIpan-3.0, a common pan-specific MHC class II prediction method including all three human MHC class II isotypes, HLA-DR, HLA-DP and HLA-DQ. Immunogenetics, 65(10): 711–724, 2013.

[45] Richèl J C Bilderbeek. tmhmm, 2019. https://github.com/richelbilderbeek/tmhmm [Accessed: 2019-03-08].

[46] Richèl J C Bilderbeek. pureseqtmr, 2020. https://github.com/richelbilderbeek/pureseqtmr [Accessed: 2020-05-19].

[47] Richèl J C Bilderbeek. netmhc2pan, 2019. https://github.com/richelbilderbeek/netmhc2pan [Accessed: 2019-03-08].

[48] Richèl J C Bilderbeek. bbbq, 2020. https://github.com/richelbilderbeek/bbbq [Accessed: 2020-09-02].

[49] Steffen Möller, Michael DR Croning, and Rolf Apweiler. Evaluation of methods for the prediction of membrane spanning regions. Bioinformatics, 17(7):646–653, 2001.

[50] Claus Lundegaard, Ole Lund, and Morten Nielsen. Prediction of epitopes using neural network based methods. Journal of immunological methods, 374(1-2):26–34, 2011.

[51] Morten Nielsen, Claus Lundegaard, Peder Worning, Sanne Lise Lauemøller, Kasper Lamberth, Søren Buus, Søren Brunak, and Ole Lund. Reliable prediction of T-cell epitopes using neural networks with novel sequence representations. Protein Science, 12(5):1007–1017, 2003.

[52] Edita Karosiene, Claus Lundegaard, Ole Lund, and Morten Nielsen. NetMHCcons: a consensus method for the major histocompatibility complex class I predictions. Immunogenetics, 64(3):177–186, 2012.

[53] Morten Nielsen, Claus Lundegaard, Peder Worning, Christina Sylvester Hvid, Kasper Lamberth, Søren Buus, Søren Brunak, and Ole Lund. Improved prediction of MHC class I and class II epitopes using a novel Gibbs sampling approach. Bioinformatics, 20(9):1388–1397, 2004.

[54] David J. Winter. rentrez: an R package for the NCBI eUtils API. The R Journal, 9:520–526, 2017.

[55] Richèl J C Bilderbeek. sprentrez, 2021. https://github.com/richelbilderbeek/sprentrez [Accessed: 2021-02-09].

[56] Lucia Musumeci, Jonathan W Arthur, Florence SG Cheung, Ashraful Hoque, Scott Lippman, and Juergen KV Reichardt. Single nucleotide differences (SNDs) in the dbSNP database may lead to errors in genotyping and haplotyping studies. Human mutation, 31(1):67–73, 2010.

[57] Ryan Hunt, Zuben E Sauna, Suresh V Ambudkar, Michael M Gottesman, and Chava Kimchi-Sarfaty. Silent (synonymous) SNPs: should we care about them? Single nucleotide polymorphisms, pages 23–39, 2009.

